# Crocodylomorph cranial shape evolution and its relationship with body size and ecology

**DOI:** 10.1101/724609

**Authors:** Pedro L. Godoy

## Abstract

Crocodylomorpha, which includes living crocodylians and their extinct relatives, has a rich fossil record, extending back for more than 200 million years. Unlike modern semi-aquatic crocodylians, extinct crocodylomorphs exhibited more varied lifestyles, ranging from marine to fully terrestrial forms. This ecological diversity was mirrored by a remarkable morphological disparity, particularly in terms of cranial morphology, which seems to be closely associated with ecological roles in the group. Here, I use geometric morphometrics to comprehensively investigate cranial shape variation and disparity in Crocodylomorpha. I quantitatively assess the relationship between cranial shape and ecology (i.e. terrestrial, aquatic, and semi-aquatic lifestyles), as well as possible allometric shape changes. I also characterise patterns of cranial shape evolution and identify regime shifts. I found a strong link between shape and size, and a significant influence of ecology on the observed shape variation. Terrestrial taxa, particularly notosuchians, have significantly higher disparity, and shifts to more longirostrine regimes are associated with large-bodied aquatic or semi-aquatic species. This demonstrates an intricate relationship between cranial shape, body size and lifestyle in crocodylomorph evolutionary history. Additionally, disparity-through-time analyses were highly sensitive to different phylogenetic hypotheses, suggesting the description of overall patterns among distinct trees. For crocodylomorphs, most results agree in an early peak during the Early Jurassic and another in the middle of the Cretaceous, followed by nearly continuous decline until today. Since only crown-group members survived through the Cenozoic, this decrease in disparity was likely the result of habitat loss, which narrowed down the range of crocodylomorph lifestyles.

## Introduction

The relationship between form and function has long been recognised (Cuvier, 1817; Russell, 1916; Lauder, 1981) and, given the phenotypic similarities generated by convergence, the incorporation of phylogenetic comparative methods has become almost imperative on analyses of evolutionary shape changes (Bookstein *et al*., 1985; Felsenstein, 1985; Harvey & Pagel, 1991; Rohlf, 2001, 2002; Losos, 2011; Monteiro, 2013). Taking this into account, several studies have examined the association between organisms’ shape and ecology in a phylogenetic context (e.g. Sidlauskas, 2008; Bhullar *et al*., 2012; Watanabe *et al*., 2019). Similarly, another widely studied and documented evolutionary phenomenon is the link between size and shape, which generates allometric shape changes (Gould, 1966; Klingenberg, 2016). Accordingly, with the current expansion of the use of geometric morphometrics techniques for analysing shape variation, studies that investigate the relationship between shape and either size or ecology (or both), while also taking a phylogenetic approach, have become increasingly common (Adams *et al*., 2004; Zelditch *et al*., 2012).

In this context, data collected from fossil organisms can yield essential information for a better comprehension of large-scale evolutionary shape changes. Among tetrapods, Crocodylomorpha represent a good system for studying large-scale phenotypic evolution, given the group’s long and rich fossil record (Bronzati *et al*., 2015; Mannion *et al*., 2015), as well as extensive recent effort to resolve major phylogenetic uncertainties (e.g. Jouve *et al*., 2006; Larsson & Sues, 2007; Young & Andrade, 2009; Young *et al*., 2010; Andrade *et al*., 2011; Clark, 2011; Brochu, 2011, 2012; Bronzati *et al*., 2012; Montefeltro *et al*., 2013; Pol *et al*., 2014; Herrera *et al*., 2015; Turner, 2015; Wilberg, 2015). Furthermore, previous studies have investigated the relationship between form and function in crocodylomorphs, particularly focusing on the link between ecological roles and skull shape (Taylor, 1987; Busbey, 1995; Brochu, 2001). Historically, the crocodylomorph skull has received substantial attention in anatomical studies (Iordansky, 1973), which might explain the preference for this part of the skeleton as the source of morphological information in most works quantitatively investigating phenotypic evolution in the group (even though some important exceptions exist; e.g. Bonnan *et al*., 2008; Chamero *et al*., 2013, 2014; Stubbs *et al*., 2013;Walmsley *et al*., 2013; Toljagić & Butler, 2013; Gold *et al*., 2014).

Previous works that use geometric morphometrics for studying crocodylomorph cranial shape have mostly focused on specific subgroups, especially crocodylians (Monteiro *et al*., 1997; Pierce *et al*., 2008; Sadleir & Makovicky, 2008; Piras *et al*., 2009, 2010, 2014; Pearcy & Wijtten, 2011; Watanabe & Slice, 2014; Okamoto *et al*., 2015; Clarac *et al*., 2016; Salas-Gismondi *et al*., 2016, 2018; Iijima, 2017; McCurry *et al*., 2017*a*; Foth *et al*., 2017; Bona *et al*., 2018; Fernandez Blanco *et al*., 2018; Morris *et al*., 2019), but also thalattosuchians (Pierce *et al*., 2009; Young *et al*., 2010) and notosuchians (Godoy *et al*., 2018). One important exception is the recent work of Wilberg (2017), that assessed cranial shape variation across Crocodyliformes (which is only slightly less inclusive than Crocodylomorpha; Irmis *et al*., 2013), sampling a large number of species. Nevertheless, the sample size of Wilberg (2017) could still be significantly increased, potentially permitting the assessment of morphospace occupation and morphological disparity among other crocodylomorph subgroups (i.e. not only crown and non-crown group species). Furthermore, Wilberg (2017) analysed patterns of cranial shape disparity through time within Crocodylomorpha, but the impact of alternative time sub-sampling methods on disparity-through-time analyses (as recently suggested by Guillerme & Cooper [2018]) was not explored, as well as that of distinct phylogenetic hypotheses. Finally, the potential influence of body size and ecological transitions on crocodylomorph cranial shape can also be quantitatively assessed with phylogenetic comparative methods (Zelditch *et al*., 2012; Klingenberg & Marugán-Lobón, 2013; Monteiro, 2013). Even though the hypothesis of a link between cranial shape and ecology (mainly feeding strategies) have been previously examined for some groups (e.g. Busbey, 1995; McHenry *et al*. 2006; Young *et al*., 2010), a wider investigation, including taxa of all crocodylomorph groups, remains to be tested, as well as the role of size in this relationship.

Here, I use geometric morphometric techniques to comprehensively analyse crocodylomorph cranial shape, by combining a previously available landmark dataset (from Wilberg [2017]) with newly digitised specimens. I quantify cranial shape variation and estimate disparity of distinct crocodylomorph subgroups, and also estimate disparity through time. This allowed me to compare my results with those of previous studies, but also to investigate the impact of a variety of alternative methods in disparity-through-time estimation. I further investigate the association between the observed shape variation and two factors: body size and ecology (=lifestyles). For that, I used disparity estimation and morphospace occupation, but also statistical and phylogenetic comparative methods. I also inferred shifts in cranial shape evolutionary regimes, using Bayesian and maximum-likelihood approaches. By doing this, I was able to characterise the patterns of crocodylomorph cranial shape evolution and to test its hypothesised relationship with ecological transitions and size.

## Material and Methods

### Sampling and data collection

Most crocodylomorph crania are taphonomically deformed, usually by dorsoventral compression, which prevents a comprehensive study from using three-dimensional (3D) data. Thus, I initially used the data published by Wilberg (2017), as this is the most phylogenetically comprehensive 2D landmark dataset to date (i.e. 131 crocodylomorph specimens, most of which were identified to species level). This dataset used only dorsal views of skulls, as this view is less susceptible to compression and taphonomic distortion across different crocodylomorph groups. Wilberg (2017) digitised four landmarks, each at a key homologous point of crocodylomorph skulls, and a semilandmark curve to represent the outline of skulls. It is worth mentioning that the dataset made available by Wilberg (2017) was slightly modified, as in its original version the semilandmark curves were artificially divided into two parts (one rostral to and another caudal to the level rostralmost point of the orbit; see Appendix A for description and position of the landmarks and further information on data collection).

To expand this dataset, I digitised landmarks for additional 86 new specimens, representing an increase of nearly 65% over the dataset of Wilberg (2017). Five specimens included in the original dataset were posteriorly removed, and the taxonomy of all specimens was updated following more recent literature (see Appendix A for further details). The final expanded dataset includes 212 specimens, representing 209 species (see Appendix B for the complete list of specimens sampled). For landmark data collection, I used tpsUTIL version 1.76 (Rohlf, 2015) to compile the images into a single .tps file, then digitising the landmarks and semilandmarks in tpsDIG2 version 2.30 (Rohlf, 2015).

### Phylogenetic framework

For the phylogenetic framework, I used a modified version of the crocodylomorph informal supertree from Godoy *et al*. (2019), with the inclusion of 20 species for which I had landmark data, but were not previously in the supertree (see Appendix A for details). The final version of the supertree includes 325 species (316 crocodylomorphs and nine non-crocodylomorph species, as outgroups). Major phylogenetic uncertainties of the group concern the relative positions of thalattosuchians (as neosuchians or sister to Crocodyliformes; Clark, 1994; Pol & Gasparini, 2009; Wilberg, 2015) and that of gavialids in relation to tomistomines and “thoracosaurs” (Gatesy *et al*., 2003; Lee & Yates, 2018). To accommodate these, three alternative phylogenetic scenarios were considered for downstream analyses, creating three topologies with variable positions of Thalattosuchia and distinct interrelationships within Crocodylia (see Appendix A for details).

Each of these topologies were then time-calibrate using the fossilised birth-death (FBD) model (Stadler, 2010; Ronquist *et al*., 2012*a*; Heath *et al*., 2014; Zhang *et al*., 2015). I used the protocol within the R package *paleotree* version 3.1.3 (Bapst, 2012), which follows recommendations within Matzke & Wright (2016), to generate an “empty” morphological matrix and perform Bayesian Markov chain Monte Carlo (MCMC) tip-dating analyses in MrBayes version 3.2.6 (Ronquist *et al*., 2012*b*). The three supertree topologies (representing alternative phylogenetic scenarios) were used as topological constraints and the uniform priors for the age of tips were set based on the occurrence dates information (i.e. obtained from the literature and the Paleobiology Database). A uniform prior was used for the root of the tree (for all three topologies), constraining it between 245 and 260 Myr ago (given that a crocodylomorph origin older than the Early Triassic is unlikely; Irmis *et al*., 2013; Turner *et al*., 2017; Ezcurra & Butler, 2018). For each alternative phylogenetic scenario, 10,000,000 generations were used in two MCMC runs (with four chains each), after which the parameters indicated that both runs converged (i.e., the Potential Scale Reduction Factor approached 1.0 and average standard deviation of split frequencies was below 0.01). In downstream analyses, for each phylogenetic scenario I used either the maximum clade credibility (MCC) trees or 10 trees randomly sampled from the posterior distribution, both after a burn-in of 25%.

### Geometric morphometric analyses

Geometric morphometric analyses were performed using the package *geomorph* (version 3.0.7; Adams & Otárola-Castillo, 2013) in R (version 3.5.1; R Core Team, 2018). Generalised Procrustes analysis (GPA) (Gower, 1975; Rohlf & Slice, 1990) was performed prior to all analyses. Semilandmarks were defined using function *define.sliders()*, and the location of sliding semilandmarks during GPA-alignment was optimised by minimizing the bending energy between the reference and target specimen (i.e. argument ProcD=FALSE within in the function *gpagen()*; see Bookstein, 1997). Subsequently, the Procrustes coordinates of specimens were used as the input data for principal component analysis (PCA; Hotelling, 1933). As the landmark data used here derives from two distinct sources (i.e. the dataset provided by Wilberg [2017] and the data collected for the present study), particular attention was paid to the potential impact of interobserver error on the cranial shape data. To quantitatively approach this issue, I used Procrustes ANOVA (i.e. linear models; Goodall, 1991; Klingenberg & McIntyre, 1998; Anderson, 2001) to compute the amount of variation caused by interobserver error (see Appendix A for further details).

To better visualise the morphospace occupation of different crocodylomorph subgroups (e.g. in PC1 versus PC2 scatterplots), I used distinct colours and convex hulls (i.e. the area inside the minimum convex polygon; Cornwell *et al*., 2006) for six taxonomic groups: Crocodylia (with “thoracosaurs”), Tethysuchia, Thalattosuchia, Notosuchia and two paraphyletic groupings of non-crocodylian neosuchians (excluding tethysuchians and thalattosuchians) and non-mesoeucrocodylian crocodylomorphs (excluding thalattosuchians). To statistically assesses the differences in the morphospace occupied by these groups, I used non-parametric multivariate analysis of variance (npMANOVA, performed with R package *RVAideMemoire*; Hervé, 2018) with all PC scores (see Appendix A for further details). To examine the influence of phylogenetic history on the observed shape variation, I calculated the phylogenetic signal (K_mult_; Bloomberg *et al*., 2003; Adams, 2014) using function *physignal()*, with Procrustes coordinates of specimens and the MCC tree of each alternative phylogenetic scenario, performing 1000 iterations.

I also divided crocodylomorph species into three categories representing distinct ecologies (or lifestyles): marine/aquatic, freshwater/semi-aquatic and terrestrial species. For that, I used information available in the literature (Mannion *et al*., 2015; Wilberg *et al*., 2019), as well as in the Paleobiology Database (see Appendix B for the lifestyles assigned to each taxon). I then visually assessed the differences in morphospace occupation using colours and convex hulls, and statistically scrutinised these differences using npMANOVA.

### Estimating disparity

For this paper, I selected the sum of variances as the disparity metric, as it seems to be more robust for measuring morphological disparity through time (see Wills *et al*., 1994; Butler *et al*., 2012; Guillerme & Cooper, 2018) and for the comparison with the results from other studies (e.g. Stubbs *et al*., 2003; Toljagić & Butler, 2013; Wilberg, 2017). Other methods were proposed to measure disparity, and often produce different results, but were not used herein (see Appendix A for further discussion).

I performed a series of sensitivity analyses for disparity-through-time estimation, given that previous work demonstrated the susceptibility of these kind of analyses to alternative methods for sub-sampling taxa (Guillerme, 2018; Guillerme & Cooper, 2018). Thus, I used different sub-sampling procedures (i.e. time binning and time-slicing methods, *sensu* Guillerme & Cooper [2018]), different numbers of time intervals (10 and 20), and multiple time-scaled phylogenetic trees (randomly sampled from the posterior distribution of tip-dating MCMC analyses) to assess the impact of these alternative approaches on the results (see Appendix A for further information on sensitivity analyses).

I also estimated disparity (=sum of variances) among different crocodylomorph subgroups (taxonomic and ecological groups), by dividing the PC scores for each species into distinct subsets. PC scores were subsequently bootstrapped 100 times and, during each bootstrap replication, the number of elements (taxa) drawn was standardised in all groups, by rarefying the data (i.e. using the minimum number of species in a subset, which was 18 and 48 elements for taxonomic and ecological subsets, respectively). Significant differences in bootstrapped median values were statistically assessed using npMANOVA with 10,000 permutations, followed by a Bonferroni correction for adjusted *p*-values (Rice, 1989, Anderson, 2001). All disparity analyses (through-time and between groups) were performed using the R package *dispRity* (Guillerme 2018).

### Correlation with body size and ecological factors

I used Procrustes ANOVA, with function *procD.allometry()* in *geomorph*, to investigate and visualise allometric changes and how much of shape variation can be explained by body size. I used log-transformed centroid size as a proxy for total body size and calculated the regression scores (Drake & Klingenberg, 2008) for plotting purposes. To further inspect and visualize this possible shape-size relationship, I regressed and plotted my cranial shape data against an independent body size dataset (a comprehensive crocodylomorph dataset of log-transformed dorsal cranial length measurements, made available by Godoy *et al*. [2019]), using ordinary least square (OLS) and phylogenetic generalized least squares (PGLS) regressions. For PGLS, I incorporated the phylogenetic information from the maximum clade credibility (MMC) trees (of each alternative phylogenetic scenario) and optimized branch length transformations between bounds with maximum-likelihood using Pagel’s λ (Pagel, 1999) (i.e., argument λ = “ML” within in the function *pgls()* of the R package *caper*; Orme *et al*. 2018).

In addition to estimating disparity for distinct ecological categories (see above), I further assessed the influence of ecological factors on crocodylomorph cranial shape by applying Procrustes ANOVA in a phylogenetic framework, using *procD.pgls()* function in *geomorph*. As for allometric Procrustes ANOVA, 10,000 permutations were performed, and I obtained the percentage of variation explained by the independent variables (body size or ecology) by dividing the sum of squares of the variable by the total sum of squares.

### Identifying regime shifts in crocodylomorph cranial shape evolution

Some studies have previously indicated difficulties and intrinsic issues of many of the presently proposed methods for automatically detecting regime shifts of phenotypic traits in phylogenies (e.g. Adams & Collyer, 2018; Bastide *et al*., 2018), many of which assume evolution under a non-uniform Ornstein-Uhlenbeck [OU] process (although methods that assume other models/processes are also available; e.g. Rabosky, 2014; Castiglione *et al*., 2017, 2019). In particular, the combination of using multivariate data (such as shape data derived from geometric morphometric methods) and fossils as tips in a time-scaled tree (i.e. a non-ultrametric tree) presents a challenge to currently proposed methods, without an appropriate solution to date (Bastide *et al*., 2018).

Accordingly, I decided to reduce my analyses to a single (univariate) trait, using only PC1 scores of taxa, since this principal component represents a significant amount of total shape variation (more than 70%) and is biologically meaningful (i.e. translating, predominantly, changes in snout length). Using only one dimension (i.e. PC1) allowed me to apply two methods for detecting regime shifts: *bayou* (Uyeda & Harmon, 2014) and *SURFACE* (Ingram & Mahler, 2013). Differently from other methods proposed (e.g. *l1ou* [Khabbazian *et al*., 2016] and *PhylogeneticEM* [Bastide *et al*., 2018]), *bayou* and *SURFACE* can deal with non-ultrametric trees, and seem to work better with univariate data (Adams & Collyer, 2018). Even though not ideal (see Polly *et al*. [2013], Uyeda *et al*. [2015], Adams & Collyer [2018], and Du [2018] for problems with dimension reduction), by comparing the outputs from both methods (i.e. looking for overall patterns of shift detection) I could visualise patterns of cranial shape variation along the crocodylomorph phylogeny, as well as to compare to body size evolutionary patterns and to ecological transitions in the group.

For *bayou*, which is a Bayesian reversible-jump approach (Uyeda & Harmon, 2014), I ran five MCMC chains of 1,000,000 generations for each of the three phylogenetic scenarios (using the MCC trees), with 30% burn-in and a conditional Poisson distribution as a prior on the number of shifts. For *SURFACE*, which is a stepwise AIC procedure (Ingram & Mahler, 2013), I used phylogenetic Bayesian information criterion (pBIC) as an alternative to AICc in the backward-phase of SURFACE (during which “convergent” regimes are identified), as the former is more conservative than AICc, generating lower rates of false positive identification of regime shifts (Ho & Ané, 2014; Khabbazian *et al*., 2016; Benson *et al*., 2018). Furthermore, given that SURFACE seems to be very sensitive to both topological conformation and branch lengths, instead of MCC trees I used 30 time-scaled crocodylomorph trees for the analyses (i.e. 10 randomly sampled trees of each alternative phylogenetic scenario), and assessed regime shift identification on all trees, looking for overall patterns (see Appendix A for more details on *SURFACE* and *bayou* analyses). *bayou* analyses were performed with R package *bayou* (Uyeda *et al*., 2018), whereas *SURFACE* analyses were performed with *surface* package (Ingram & Mahler, 2013). Implementation of pBIC functions in the backward-phase of *SURFACE* model fits used scripts made available by Benson *et al*. (2018). See Appendix C for an R script with the all the analyses performed here (i.e. geometric morphometric, disparity, *SURFACE*, and *bayou* analyses), as well as landmark data and phylogenetic trees.

## Results

### Cranial shape in different crocodylomorph subgroups

Procrustes ANOVA results show that interobserver error accounts for only 1.6% of total shape variation (Appendix A Table S1), allowing further analyses using the expanded dataset. The aspects of morphology represented by PC1 and PC2 (Fig. 1) are equivalent to those found by Wilberg (2017), with PC1 (71.89% of the variation) mostly describing variation in snout length and PC2 (8.6%) changes in the quadrate condyle and the position of the orbit in relation to the lateral outline of the skull (see Appendix A Fig. S3 for variation in all PCs).

**Figure 1.**
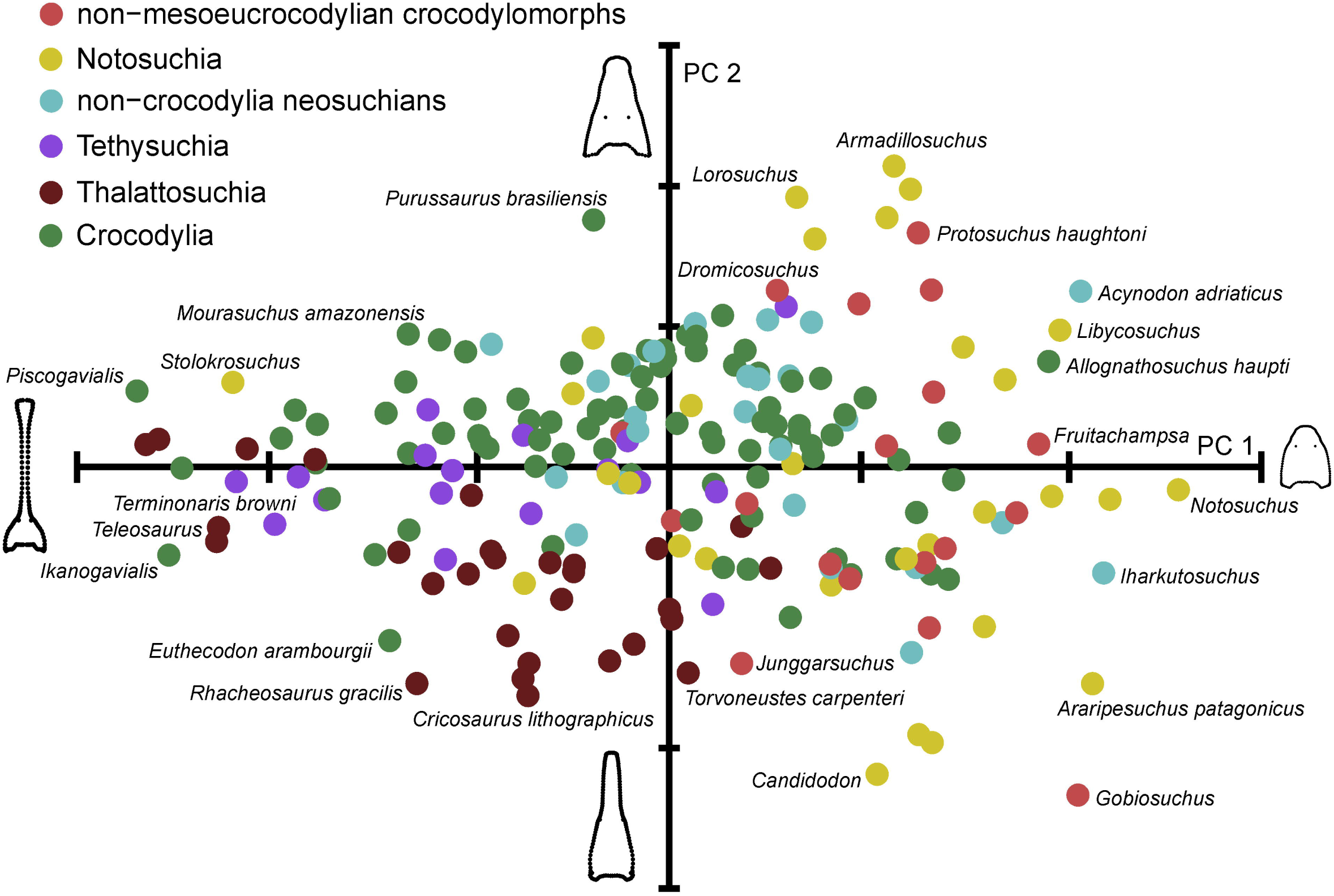
Morphospace plot (PC1 versus PC2) of crocodylomorph cranial shape variation, using the expanded dataset, after removing the effect of interobserver error (by standardising the landmark digitising protocol). Different colours represent distinct crocodylomorph subgroups: non-mesoeucrocodylian crocodylomorphs (excluding thalattosuchians), Notosuchia, non-crocodylian neosuchians (excluding tethysuchians and thalattosuchians), Tethysuchia, Thalattosuchia, and Crocodylia. PC1 and PC2 represent, respectively, 71.89% and 8.6% of total shape variation.

Comparisons of the morphospaces occupied by different crocodylomorph subgroups (Fig 2a, b, and Appendix A Fig. S4) reveal a wide distribution of members of the crown-group (crocodylians) and notosuchians. Crocodylians exhibit almost the entire range of morphological variation described by PC1 (i.e. long versus short snouts), whereas most notosuchians occupy the region of short rostra (although the presence of *Stolokrosuchus lapparenti* in the analysis expands the morphospace occupation of the group towards the “longirostrine region”; Fig 2a, b, and Appendix A Fig. S4). Pairwise statistical assessment using npMANOVA (Appendix A Table S2) reinforces the apparently disparate cranial morphology of these two groups, as it shows that their morphospaces are significantly different to one another (*p* = 0.0015), and also from most of the groups tested (see Appendix A for further description of morphospace occupation in other crocodylomorph subgroups).

**Figure 2.**
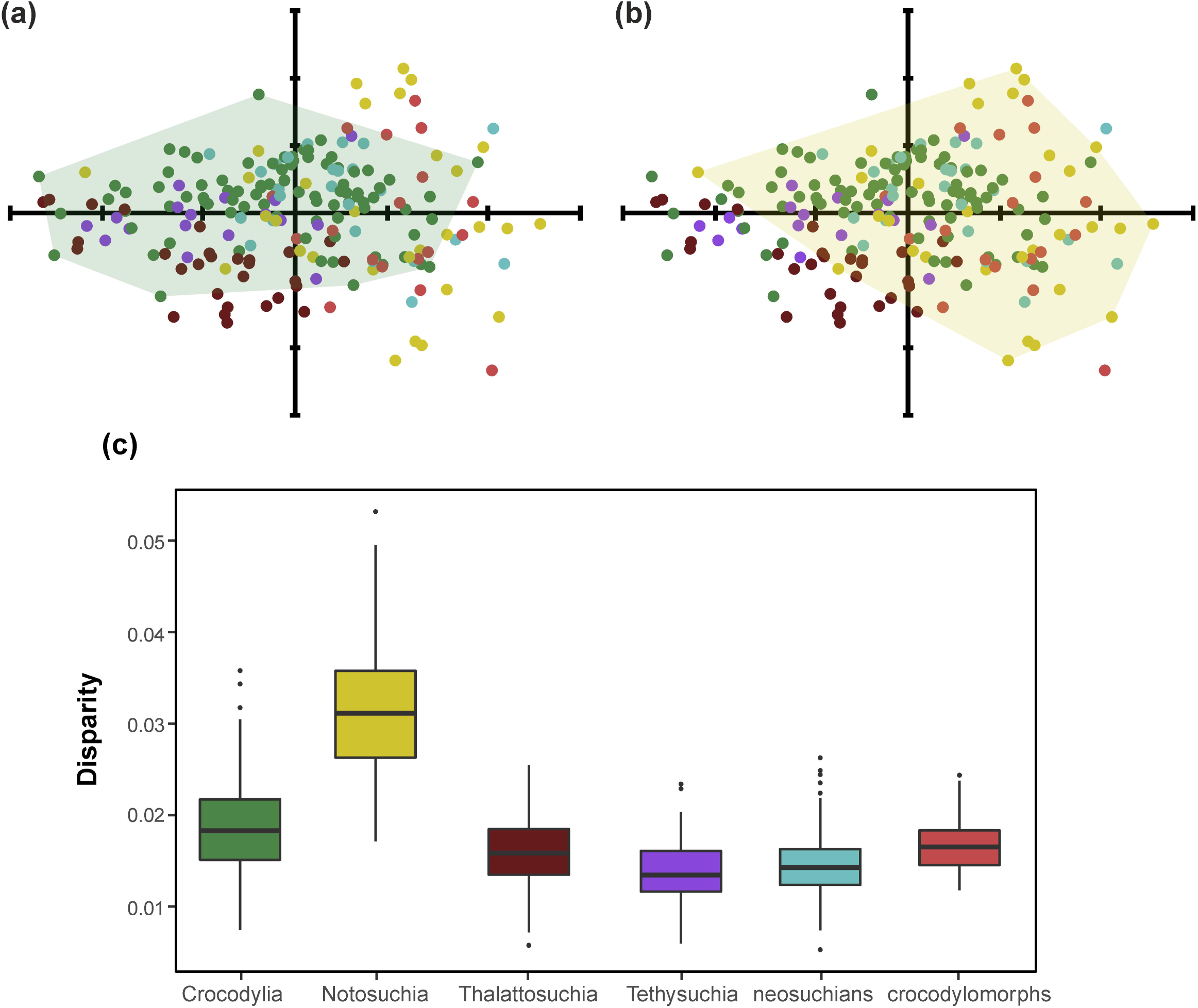
Morphospace occupation and cranial shape disparity (sum of variances) of different crocodylomorph subgroups. (a) Morphospace occupied by members of the crown-group (Crocodylia). (b) Morphospace occupied by notosuchians. (c) cranial shape disparity of species divided into six taxonomic subsets (“neosuchians” represent non-crocodylian neosuchians, without tethysuchians and thalattosuchians; “crocodylomorphs” represent non-mesoeucrocodylian crocodylomorphs, without thalattosuchians). PC scores of specimens were bootstrapped and rarefied for disparity calculation.

Cranial shape disparity estimated for different crocodylomorph subgroups revealed that Notosuchia has the highest cranial shape disparity among all groups assessed (Fig. 2c). Crocodylia exhibits a smaller disparity, although slightly higher than the other four groups (Tethysuchia, Thalattosuchia, non-crocodylian neosuchians, and non-mesoeucrocodylian crocodylomorphs), which have comparable median values. Pairwise comparisons (Appendix A Table S3) show that disparity in both Notosuchia and Crocodylia is significantly different from that in all other groups analysed, whereas some other groups have statistically equivalent disparities (e.g. thalattosuchians and non-mesoeucrocodylian crocodylomorphs, as well as tethysuchians and non-crocodylian eusuchians). Similar results were recovered when fewer subsets of taxa were analysed (i.e. Notosuchia, Neosuchia, Thalattosuchia, non-mesoeucrocodylian crocodylomorphs), with notosuchian disparity still higher and significantly different from the other groups (Appendix A Fig. S5 and Table S4).

Even though some crocodylomorph subgroups exhibit morphospaces that are significantly distinct from other groups, the relatively weak to moderate phylogenetic signal estimated for the data (K_mult_ values varied between 0.0866 and 0.2398 with different phylogenetic scenarios; *p* < 0.05) suggest no strong degree of phylogenetic structure in patterns of cranial shape variation among taxa. These results are consistent with the visual representation of phylogenetic information is incorporated into tangent space (i.e. phylomorphospace plots of PC1 against PC2; see Appendix A Fig. S6), with the multiple intersections of branches.

### Disparity through time

In general, significant impacts on disparity-through-time analyses were observed when distinct tree topologies were used (see Appendix D for plots of all disparity-through-time analyses). Similarly, distinct time sub-sampling methods (i.e. number of time intervals used and the use of time bins or time-slices) also impacted in disparity estimation. The only exception to this was when assuming different evolutionary models (i.e., punctuated or gradual model) for time-slicing method, which produced almost identical results (Appendix A Figs. S7, S8, and S9; see Appendix D for all plots). Comparisons between the 10 trees within a same phylogenetic scenario show some dissimilarities in the pattern of disparity through time, and these differences are usually more marked when a greater number of time intervals is used (using either the time binning or the time-slicing methods). For example, analyses using distinct trees within a same phylogenetic scenario (i.e. Thalattosuchia sister to Crocodyliformes and gavialids within Gavialoidea) disagree on the timing and magnitude of a disparity peak during the early evolution of the group (Fig. 3). Whereas some trees show this peak beginning prior to the Triassic-Jurassic (T–J) boundary, other trees yield a later start, only after the boundary. Other differences include whether there is an increase or a decrease in disparity from the middle of the Neogene (Eocene) to the Recent, as well as if a peak observed during the Early Cretaceous corresponds to the highest disparity seen in the group’s entire evolutionary history (Fig. 3). Similarly, the use of alternative phylogenetic scenarios also impacted on disparity curves. For example, different positions of Thalattosuchia had greatest impact on disparity estimation during the Jurassic, since this corresponds to the age range of thalattosuchians.

**Figure 3.**
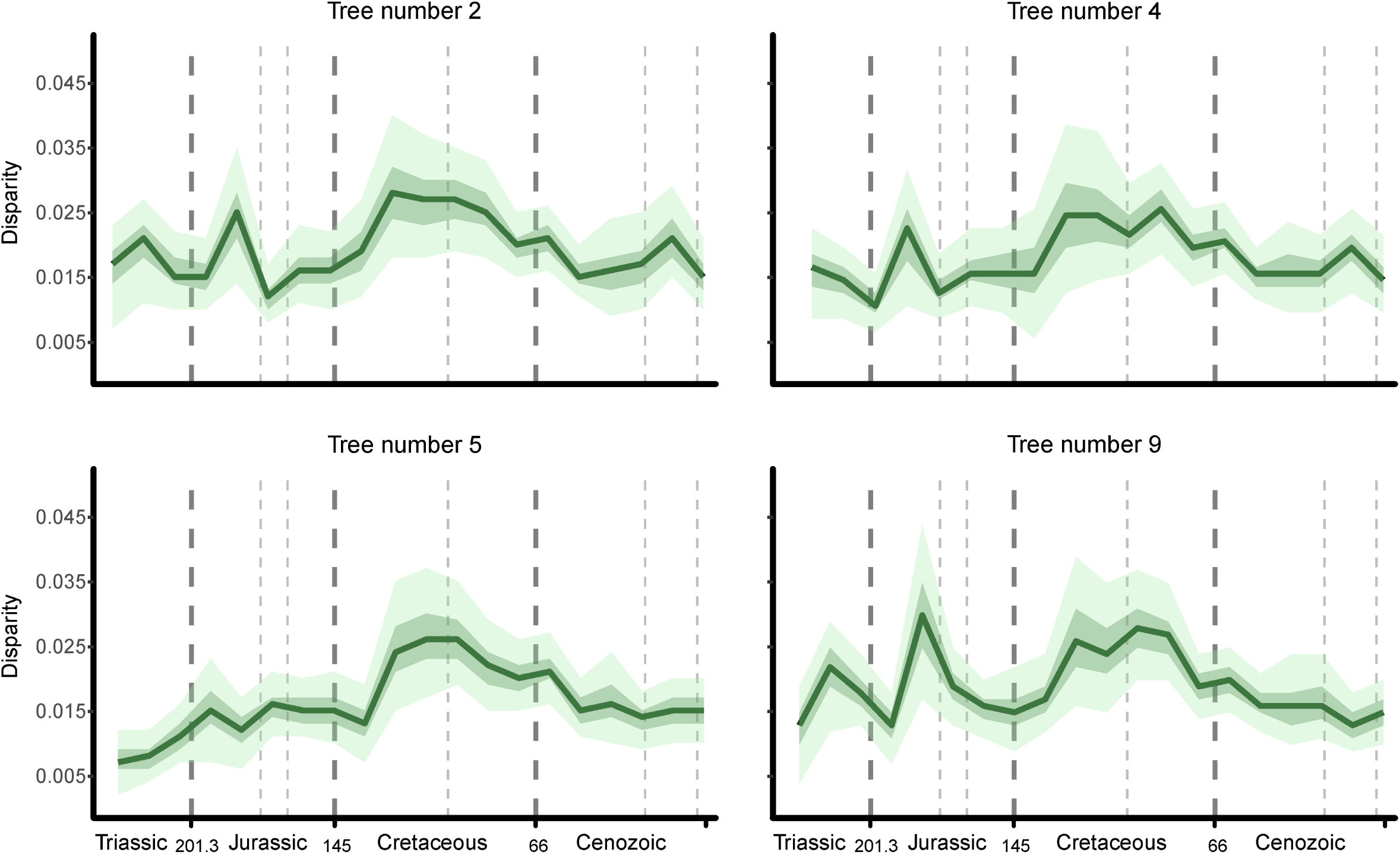
Crocodylomorph cranial shape disparity (=sum of variances) through time. Each disparity curve used a different time-scaled trees for disparity-through-time calculation. All trees share the same phylogenetic position of thalattosuchians (as sister to Crocodyliformes) and gavialids (within Gavialoidea) and used the same time sub-sampling method (time binning method, with 20 equal-length time bins). Discrepancies between results come from differences in branch lengths among trees, which reflect taxa temporal uncertainties. Light and dark green shades represent, respectively, 75% and 97.5% confidence intervals from 1,000 bootstrapping replicates.

In general, disparity-through-time analyses using more time intervals (either time bins or time slices) reconstruct more nuanced changes in disparity, even though they also often have larger confidence intervals, due to less taxa being included in each time interval (Appendix A Figs. S7, S8, and S9). When comparing different time sub-sampling methods, similar differences are observed to those seen when different tree topologies are compared (i.e. variation in the timing and magnitude of disparity peaks). For example, the magnitude estimated for the peak seen at the end of the Early Cretaceous was usually greater when using the time-slicing method.

Despite these dissimilarities arising from different time-scaled trees and time sub-sampling methods, most analyses seem to agree on some overall patterns of crocodylomorph cranial shape disparity through time (Fig. 3, Appendix A Figs. S7, S8, and S9). An early peak in disparity is frequently observed, most often during the Early Jurassic (although sometimes even prior to the Triassic-Jurassic boundary). Following a sharp decrease during the Middle Jurassic, disparity undergoes a continuous increase until the middle of the Cretaceous (Aptian–Albian), when maximum disparity is reached in most analyses. Subsequently, a near constant decline is observed during the Late Cretaceous and the Palaeocene, with analyses only disagreeing whether it continues until the Recent or ceases during the Eocene. In these latter cases (more frequently seen in analyses using the time-slicing method), a sharp increase in disparity is seen in the Eocene, but is frequently followed by an equally sharped decrease until the Recent.

These overall patterns resemble those found by Wilberg (2017) in that a clear peak is observed in the Cretaceous, followed by a nearly continuous decline towards the Recent. Some differences, however, are also noted. Since Wilberg (2017) used different disparity metrics in his analyses, my comparisons between my results and his focus on the results of variance-based disparities. The first discrepancy arises from the fact that Wilberg (2017) restricted his study to Crocodyliformes (with the exception of thalattosuchians) and did not include any Late Triassic species, resulting in an absence of information about crocodylomorph disparity prior to the Jurassic. When using stratigraphic intervals as time bins for his disparity-through-time analyses (resulting in 36 time bins), Wilberg (2017) found two significant disparity peaks during the Jurassic (one in the Pliensbachian and another in the Aalenian–Bajocian), whereas in most of my analyses a single Jurassic peak was estimated, usually occurring from the Sinemurian to the Toarcian (Fig. 3, Appendix A Figs. S7, S8, and S9). The timing of the disparity peak in the Cretaceous is another divergence between the two studies, with the analyses performed by Wilberg (2017) indicating a Late Cretaceous peak (Cenomanian), whereas most of my analyses show a slightly earlier peak (Barremian– Albian). Finally, another difference was found in the pattern of disparity from the Eocene to the Recent. Whereas many of my analyses (particularly when using the time-slicing method) indicate a disparity increase starting in the Eocene, this increase is not identified by the variance-based analyses in Wilberg (2017).

### Allometric changes in cranial shape

Body size (=centroid size) has a significant (*p* < 0.005) effect on crocodylomorph cranial shape, representing nearly 35% of the total observed variation (this is increased to more than 45% when only PC1 is considered; Table 1). This relationship can be visualised in a shape versus size plot, using the regressions scores (which shape variation represented is very similar to that of PC1) and log-transformed centroid size (Fig. 4b). Mapping PC1 (which represents more than 70% of all observed shape variation) into crocodylomorph phylogeny indicates that many of the largest taxa (such as some thalattosuchians and tethysuchians) also exhibit PC1 values associated to longer rostra, whereas most of the predominantly small-bodied notosuchians show PC1 values related to shorter snouts (Fig. 4a). Further examination of this relationship using an independent body size dataset (from Godoy *et al*. 2019) provided very similar results, with a significant correlation between shape and size (using untransformed and phylogenetically corrected data; Appendix A Table S5 and Fig. S10). These results indicate that body size is a strong predictor of cranial shape in the group. Consequently, the morphospace occupation of distinct crocodylomorph subgroups using “allometry-free” shape data (i.e. from size-adjusted residuals, Fig. 4c) reveal different patterns from that of uncorrected data (Fig. 1 and Appendix A Fig. S4), even though some general patterns can still be recognised. For example, crocodylian morphospace is comparatively more restricted, without exploring the region of extreme longirostrines (which is mainly dominated by thalattosuchians), whereas tethysuchians are more widespread, expanding their morphospace to that of more short-snouted taxa.

**Figure 4.**
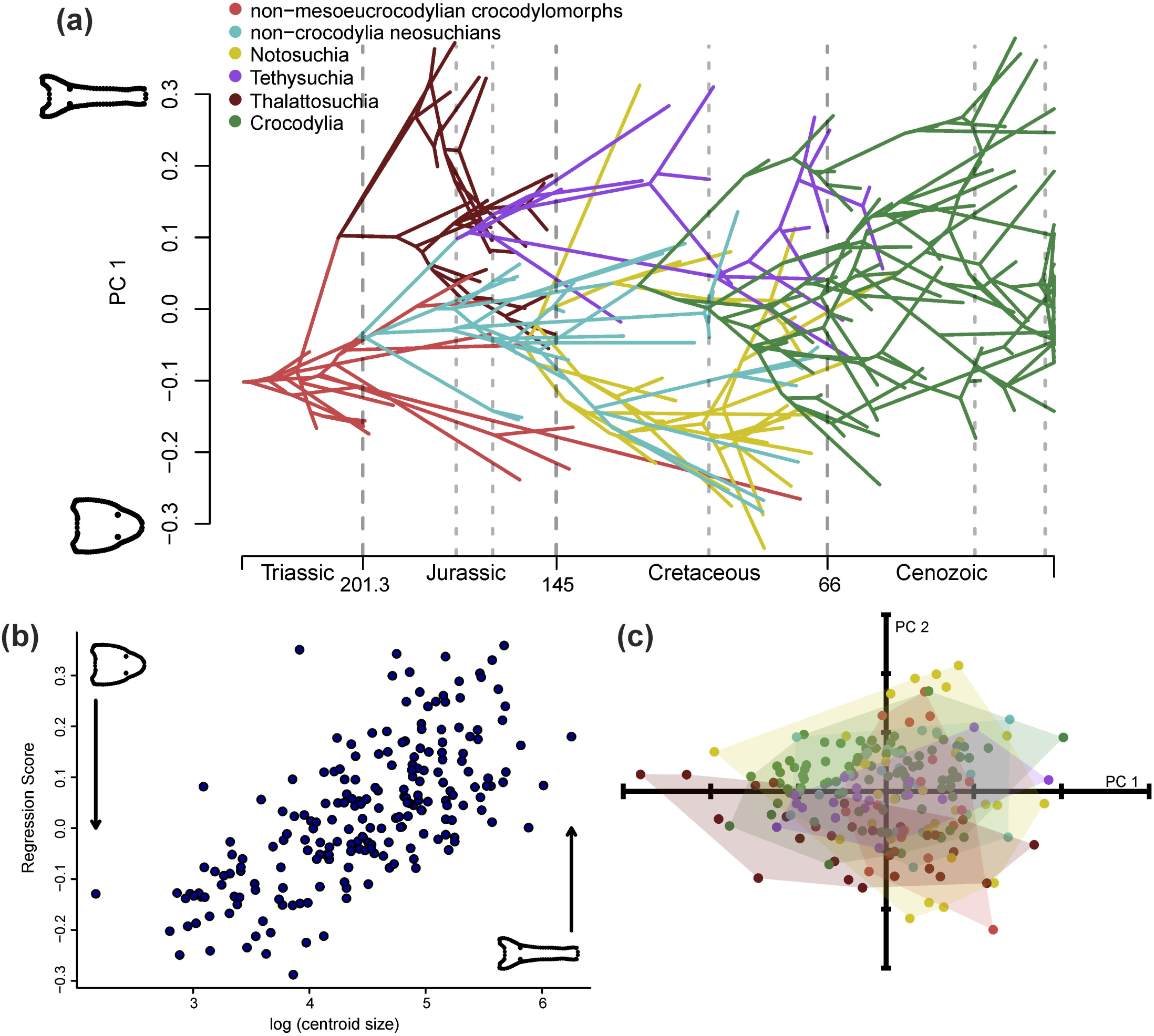
(a) Phenogram mapping crocodylomorph cranial shape (PC1 scores) onto the group’s phylogeny (topology from the MCC tree with Thalattosuchia sister to Crocodyliformes and gavialids within Gavialoidea). Different colours represent distinct mono- and paraphyletic crocodylomorph subgroups. (b) The relationship between cranial shape (regression scores) and body size (log-transformed centroid size) in crocodylomorphs. (c) PC1 versus PC2 plot showing crocodylomorph subgroups morphospace occupation using “allometry-free” shape data (i.e. from size-adjusted residuals). Colour key as in (a).

**Table 1.** Procrustes ANOVA results investigating the amount of variation in shape data explained by body size (=centroid size). **SS,** sum of squares after 10,000 permutations; **MS,** mean squares; **% of variation,** obtained by dividing the sum of squares of the independent variable (centroid size) by the total sum of squares; **F,** F-statistic; **p,** *p*-value. *Significant at alpha = 0.05.

| Total shape variation (all PCs) |  |  |  |  |  |
| --- | --- | --- | --- | --- | --- |
| Effect | SS | MS | % of variation | F | p |
| Centroid size | 1.8246 | 1.82461 | 33.70073 | 105.22 | 0.0001* |
| Residuals | 3.5895 | 0.01734 |  |  |  |
| Total | 5.4142 |  |  |  |  |
| Shape variation represented by PC1 |  |  |  |  |  |
| Effect | SS | MS | % of variation | F | p |
| Centroid size | 1.8097 | 1.80967 | 46.49398 | 179.87 | 0.0001* |
| Residuals | 2.0826 | 0.01006 |  |  |  |
| Total | 3.8923 |  |  |  |  |

### Cranial shape and ecology

Procrustes ANOVA results show a significant, although small (5%, *p* < 0.05), influence of ecology (=lifestyles) on crocodylomorph cranial shape (Table 2). Consistently, npMANOVA results indicate that all three ecological categories have significantly different morphospace occupation (i.e. *p* < 0.001). The PC1 versus PC2 scatterplot (Fig. 6a) reveal that most terrestrial taxa are restricted to the region of short-snouted skulls along PC1 axis, although they have a wider distribution along PC2 axis. Conversely, aquatic crocodylomorphs, represented by some of the most extreme longirostrine forms (such as the gavialid *Ikanogavialis gameroi*), are mainly confined to the region of longirostrine. Semi-aquatic species are more widespread along the PC1 axis (Fig. 6a), even though their distribution along the PC2 axis seems to be similar to that observed for aquatic forms. In terms of disparity (=sum of variances), terrestrial crocodylomorphs show significantly higher disparity than the other two categories, whereas aquatic and semi-aquatic species exhibit similar median disparity (Fig. 6b and Appendix A Table S6).

**Figure 5.**
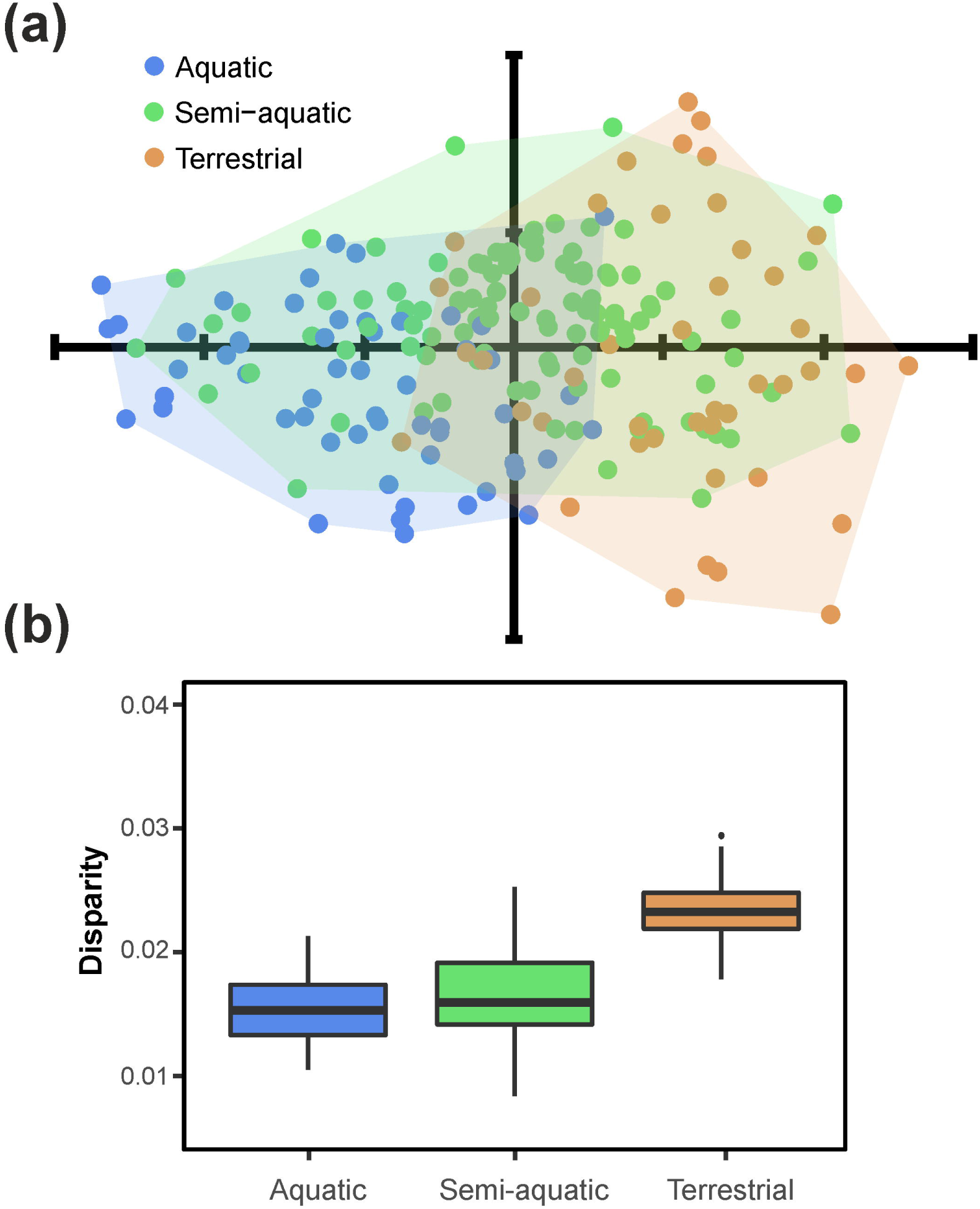
(a) Morphospace occupation of crocodylomorphs divided into three ecological categories: aquatic/marine (n = 54), semi-aquatic/freshwater (n = 107) and terrestrial (n = 48). (b) Cranial shape disparity (=sum of variances) with crocodylomorphs subdivided into the same three categories. PC scores of specimens were bootstrapped and rarefied for disparity calculation.

**Figure 6.**
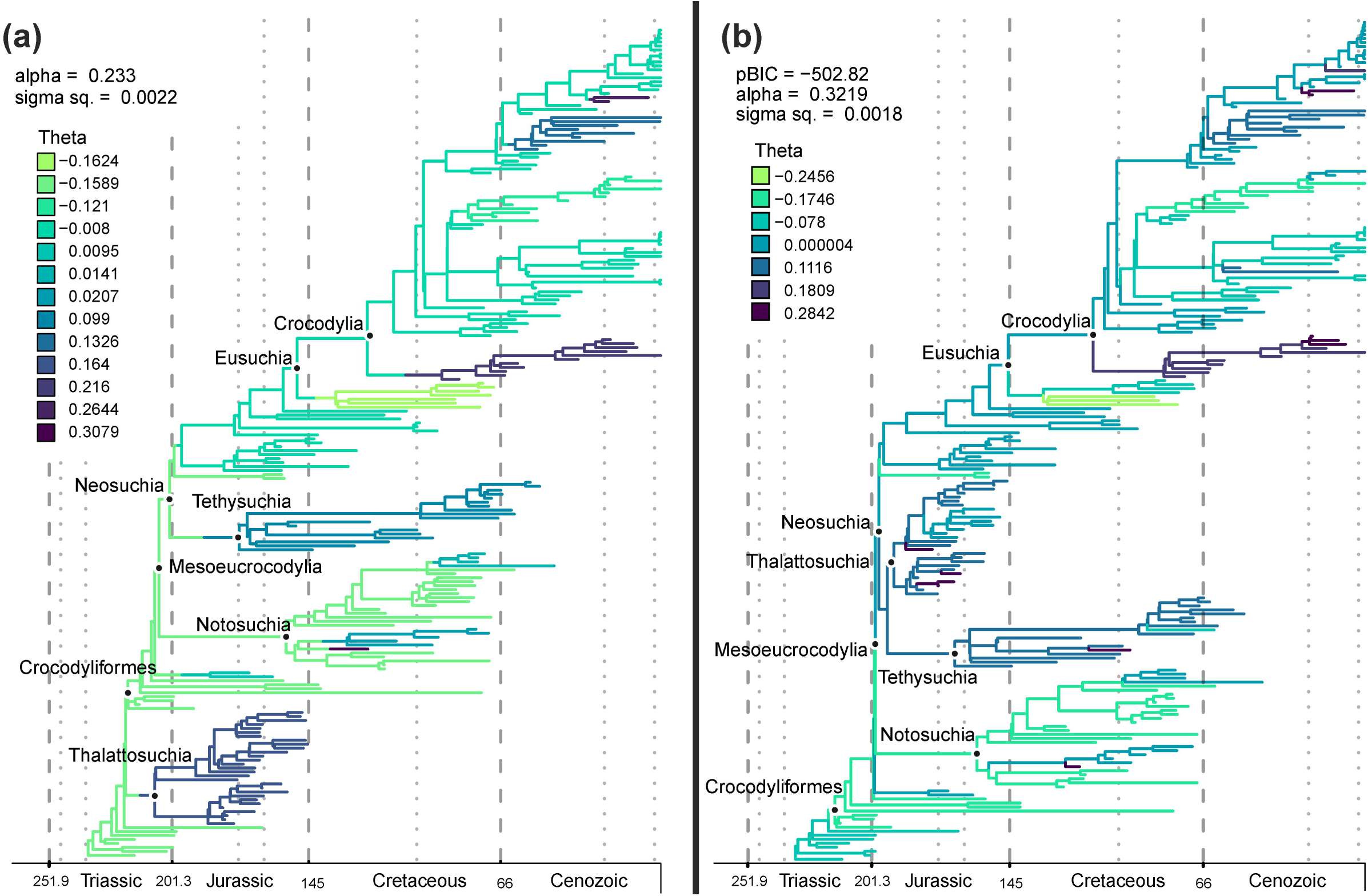
Plots with results of bayou and SURFACE analyses using PC1 scores (as a proxy for skull shape) for identifying cranial shape regime shifts in crocodylomorph evolutionary history. Branches are coloured according to different values of regime trait optima/theta (θ), with lighter colours associated to regimes of shorter snouts and darker colours representing regimes of more longirostrine taxa. (a) Results from bayou analysis using the MCC tree with Thalattosuchia sister to Crocodyliformes and gavialids within Gavialoidea. Parameter estimates alpha (α, strength of attraction) and sigma square (σ2, rate parameter) are mean values from all MCMC analyses with this tree topology (after 30% burn-in). Only theta (θ) values with posterior probability higher than 0.5 are shown. (b) Results from SURFACE analyses using tree number 10 with Thalattosuchia within Neosuchia and gavialids within Gavialoidea. Likelihood information criterion (pBIC) and parameter estimates values shown only for this tree topology. Theta values shown are those of “convergent” regimes.

**Table 2.** Procrustes ANOVA results investigating the amount of variation in shape data explained by ecology (=lifestyle). **SS,** sum of squares after 10,000 permutations; **MS,** mean squares; **% of variation,** obtained by dividing the sum of squares of the independent variable (lifestyle) by the total sum of squares; **F,** F-statistic; **p,** *p*-value. *Significant at alpha = 0.05.

| Total shape variation (all PCs) |  |  |  |  |  |
| --- | --- | --- | --- | --- | --- |
| Effect | SS | MS | % of variation | F | p |
| Lifestyle | 0.007586 | 0.0037932 | 3.098517 | 3.2935 | 0.0079* |
| Residuals | 0.237256 | 0.0011517 |  |  |  |
| Total | 0.244842 |  |  |  |  |

### SURFACE *and* bayou *analyses*

Despite some specific discrepancies (see Appendix E for plots of all *SURFACE* and *bayou* results), the results of both *SURFACE* and *bayou* analyses show similar patterns of crocodylomorph cranial shape evolution, as a clear consistency is observed in regime shifts detection across different methods (Fig. 6). In general, regime shifts are associated with ecological transitions and body size regime shifts in crocodylomorphs, which were previously presented by other studies (Wilberg *et al*., 2019; Godoy *et al*., 2019). In particular, regime shifts to longer snouts (i.e. PC1 scores trait optima values, θ, > 0.1) are frequently detected in groups of aquatic or semi-aquatic species, which are usually large-bodied animals, such as thalattosuchians, tethysuchians, gavialids and “thoracosaurs”. The opposite is commonly true for terrestrial taxa, mostly associated with regimes of short rostra (θ < −0.1), even though some exceptions exist (such as the large-bodied terrestrial sebecosuchians). The ancestral cranial shape regime was frequently associated to shorter snouts (with values of ancestral trait optimum, Z_0_, ranging from −0.16 to −0.07), which is consistent with some of the oldest known crocodylomorph taxa, such as “sphenosuchians” and protosuchids.

The use of different phylogenetic scenarios did not cause significant impacts on these overall results, with a consistent association between trait optima values and crocodylomorph subgroups across different analyses. It is worth mentioning, however, that some *SURFACE* analyses, more frequently those with Thalattosuchia placed outside Crocodyliformes, exhibited significantly simpler model fits (with much fewer regime shifts and usually unrealized low or high values of theta). This could be the results of the inability of the *SURFACE* algorithm to deal with certain tree topologies and branch lengths (similar to the suboptimal model fits identified and demonstrated by Benson *et al*., 2018), stressing the importance of using multiple time-scaled trees for *SURFACE* analyses.

In general, *SURFACE* analyses identified more regime shifts (usually more than 15 shifts) than *bayou*, which found less than 15 shifts with highly supported posterior probabilities (i.e. signal-to-noise ratio ≫ 1; Cressler *et al*., 2015; Smaers *et al*., 2017) in all three phylogenetic scenarios. It is worth mentioning that the backward-phase of *SURFACE* lumps together regimes with similar values of θ, creating “convergent” regimes (Ingram & Mahler, 2013), which were less than 10 in most of my *SURFACE* model fits. Even though slightly different number of regime shifts were found for *SURFACE* and *bayou* analyses, the magnitudes of θ values associated to different crocodylomorph subgroups were consistent across methods. Additionally, the magnitude of alpha (α, the strength of attraction) and sigma square (σ^2^, the rate of stochastic evolution) was also very similar across most of the better supported *SURFACE* analyses and all three *bayou* analyses (see Appendix E for all plots), although some *SURFACE* analyses displayed unrealistic values (i.e. extremely high or low values), which could also be caused by unsuccessful (suboptimal) model fits.

## Discussion

### Crocodylomorph snouts and feeding ecology

Most of the shape variation in crocodylomorph skulls is represented by changes in the snout, particularly in its length and width (Fig. 1). This is consistent with what was found in previous geometric morphometric studies (e.g. Pierce *et al*., 2008; Sadleir & Makovicky, 2008; Pierce *et al*., 2009; Piras *et al*., 2009; Young *et al*., 2010; Foth *et al*., 2017; Wilberg, 2017; Godoy *et al*., 2018), indicating that this region of the skull has the highest morphological variation. High variability in crocodylomorph snout length has long been acknowledged (Langston, 1973; Busbey, 1995; Brochu, 2001), even leading early taxonomists (e.g. Lydekker, 1888) to erroneously classify crocodylomorphs into different groups (Pierce *et al*., 2008). More recently, however, cladistic studies (e.g. Clark, 1994; Jouve *et al*., 2006; Pol & Gasparini, 2009; Wilberg, 2015) have suggested that convergences of crocodylomorph snouts during their evolutionary history have created a “longirostrine problem”, in which clades that are not necessarily closely related tend to be grouped together in phylogenetic analyses. In this context, the high plasticity of crocodylomorph snouts could explain the weak to moderate phylogenetic signal found for my dataset, as well as datasets of other crocodylomorph subgroups (such as crocodylians and thalattosuchians; Pierce *et al*., 2008, 2009).

Snout length can provide useful insights into ecological specializations (Taylor, 1987; Busbey, 1995; Brochu, 2001; McHenry *et al*., 2006; Pierce *et al*., 2008; Walmsley *et al*., 2013) and recent examinations of crocodylomorph cranial functional morphology further indicate strong ecological selective pressures on the snout, particularly those arising from feeding behaviour (McHenry *et al*., 2006; Gignac & O’Brien, 2016; McCurry *et al*., 2017*a*; Ballel *et al*., 2019; Gignac *et al*., 2019). Longer snouts are traditionally associated with a piscivorous diet, since the tip of the snout moves faster through water, facilitating the capture of small prey such as fish (Thorbjarnarson, 1990; McHenry *et al*., 2006; Walmsley *et al*., 2013; McCurry *et al*., 2017*b*). Thus, the widespread presence of longirostry in different crocodylomorph subgroups is presumably related to the numerous transitions to aquatic and semi-aquatic lifestyles during crocodylomorph evolutionary history (Wilberg *et al*., 2019), which are more directly connected to piscivory, highlighting the influence of ecology on the group’s cranial shape evolution.

Furthermore, snout width also has important biomechanical implications, such as impacting on hydrodynamic pressure drag (e.g. longirostrine animals compensate the higher pressure from dragging with narrower snouts; McHenry *et al*., 2006; Walmsley *et al*., 2013). Similarly, other regions of the crocodylomorph skull that vary significantly also have important implications for biomechanics and feeding strategies, such as the changes in quadrate condyle width, which are presumably associated with the craniomandibular joint (Kley *et al*., 2010; Stubbs *et al*., 2013; Ősi, 2014), even though relatively less variation is observed in these regions when compared to the snout.

### Cranial shape and size linked to ecology

We can comprehend crocodylomorph cranial shape evolution within the concept of a Simpsonian Adaptive Landscape (Simpson, 1944; 1953), which is convenient for characterizing macroevolutionary changes, since it includes ideas such as adaptive zones invasion and quantum evolution (Stanley, 1973; Hansen, 1997; 2012). This is consistent with the methodological approach used here for characterising cranial shape evolution (i.e. *bayou* and *SURFACE* methods), which assumes evolution under an OU process (even though the fit of alternative evolutionary models, such as those under Brownian motion, was not investigated here). Within the paradigm of adaptive landscapes, the different regimes of non-uniform OU models (such as *bayou* and *SURFACE*) can be interpreted as adaptive zone (Mahler & Ingram, 2014; Uyeda & Harmon, 2014). Accordingly, taking into account the selective pressures associated to these adaptive zones, shifts between macroevolutionary regimes can possibly drive large-scale patterns of phenotypic evolution.

In crocodylomorphs, the clear relationship between ecology and cranial shape and size is evidenced by the significant effects of size and lifestyle on cranial shape demonstrated here (Tables 1 and 2; Figs. 4b and 5; Appendix A Table S6 and Fig. S10). Furthermore, the evolutionary patterns of cranial shape, which were characterised here by the *bayou* and *SURFACE* results (Fig. 6), display similarities with those of body size (analysed by Godoy *et al*. [2019]) as well as with ecological transitions (demonstrated by Wilberg *et al*. [2019]). For example, shifts to regime of more longirostrine skulls are usually associated to shifts to larger-sized regimes and transitions to aquatic or semi-aquatic lifestyles. Previous studies investigating a link between larger body sizes and a more aquatic lifestyle have focused on mammals (e.g. Downhower & Blumer, 1988; Heim *et al*., 2015; Gearty *et al*., 2018), but a similar pattern was also documented for crocodylomorphs (Godoy *et al*., 2019). Within the concept of adaptive landscapes, this intricate relationship between cranial morphology, body size and ecology could be related to adaptations to an aquatic life, with selective pressures originated from intrinsic (e.g. physiological constraints associated aquatic life) and/or extrinsic factors (e.g. resources availability, such as a predominance of fish as possible preys).

The association between cranial shape and diet can also provide insights on the higher disparity seen in terrestrial taxa (Fig. 5b). Although aquatic and semi-aquatic species also explore distinct feeding strategies other than piscivory (such as durophagy; Ősi, 2014; Melstrom & Irmis, 2019), a higher variability is exhibited by terrestrial crocodylomorphs, with strategies such as herbivory, omnivory, insectivory and hypercarnivory (Ősi, 2014; Godoy *et al*., 2018; Melstrom & Irmis, 2019). The greatest contribution to this higher disparity seen in terrestrial crocodylomorphs comes from notosuchians, most of which were terrestrials and displayed exceptionally high cranial disparity (Fig. 2c), mirroring their rich fossil record (Mannion *et al*., 2015; Pol & Leardi, 2015) as well as their high morphological and body size disparities (Stubbs *et al*., 2013; Wilberg, 2017; Godoy *et al*., 2019). The drivers of such remarkable taxic diversity and morphological disparity in the group are only poorly explored, but some hint can be provided by their occurrence temporal and geographically constrained, since most notosuchians were confined to the Cretaceous of Gondwana (Mannion *et al*., 2015; Pol & Leardi, 2015), with specific environmental conditions (hot and arid climate; Carvalho *et al*., 2010). Indeed, Godoy *et al*. (2019) found evidence for more relaxed modes of body size evolution in the group, which could also be the case for other phenotypic aspects.

Apart from notosuchians, other crocodylomorph subgroups contribute to the higher disparity of terrestrial forms, mainly non-mesoeucrocodylian crocodylomorphs (such as protosuchids, gobiosuchids and shartegosuchoids; Pol & Norell, 2004; Clark, 2011; Irmis *et al*., 2013; Buscalioni, 2017; Dollman *et al*., 2018), for which a series of cranial specialisations have been previously reported (Buscalioni, 2017; Dollman *et al*., 2018). Among these, modifications related to brachycephaly (e.g. snout length reduction, rounded neurocranial shape, dorsal rotation of the mandibles, mandibular asymmetry, and tooth loss and/or orientation change; Buscalioni, 2017) are possibly associated with feeding behaviour and might represent the result of ecological selective pressures.

### Cranial shape through time

Overall, disparity-through-time results were highly sensitive to changes in the time sub-sampling method and in the phylogenetic hypothesis used (Figs. 3, Appendix A Figs. S7, S8, and S9). The considerable variation seen in these results has multiple causes. First, distinct time-scaled trees vary in assuming different stratigraphic dates for the occurrences of individual taxa (reflecting the uncertainties in the stratigraphic occurrences of most taxa used in these analyses, with many taxa known from point occurrences but with stratigraphic uncertainty often spanning two or more stages), as well as in different resolutions for polytomies (which were randomly resolved in each tree). For similar reasons, although not tested here, it is very likely that alternative time-scaling methods (e.g. *a posteriori* time-scaling approaches *sensu* Lloyd *et al*., 2016) would also impact on disparity-through-time estimation (see Bapst, 2014 for further discussion). Furthermore, as distinct trees were used for estimating the ancestral states (i.e. landmark coordinates of hypothetical ancestors), they presumably produce distinct PC scores for ancestors, which were subsequently used in disparity estimation. Similarly, distinct approaches to estimate ancestral states could also potentially impact on the results (see Ekman *et al*., 2008; Slater *et al*., 2012). Finally, the use of distinct time sub-sampling methods, as well as different numbers of time intervals (either time bins or time slices), results in different taxa being sampled in each time interval, since the rates of sedimentation (and fossilisation) are uneven in space and time (Butler *et al*., 2012; Guillerme & Cooper, 2018).

These results shed light on the importance of using multiple time sub-sampling methods for these analyses (as previously highlighted by Guillerme & Cooper, 2018), but also multiple phylogenetic hypotheses, especially for groups with major uncertainties in stratigraphic occurrence dates and phylogenetic relationships. However, many previous studies have ignored this issue, often presenting results based on one time-scaled phylogeny and one time binning approach (e.g. Brusatte *et al*., 2008*a*, *b*; Stubbs *et al*., 2013; Foth & Joyce, 2016). In fact, the discrepancies noted between the results presented here and in Wilberg (2017) could at least partially be explained by the use a single tree in the latter study (as well as by the different sample sizes and time sub-sampling methods used). Accordingly, rather than using a single analysis, perhaps a better way to report the results might be by describing shared patterns among multiple outputs, as done here.

Regarding the overall disparity through time results, the peaks and declines observed are presumably associated to the appearance and extinction of distinct crocodylomorph subgroups, such as thalattosuchians in the Jurassic and notosuchians in the Cretaceous, as already pointed out by previous studies (Stubbs *et al*., 2013; Wilberg, 2017). Some of these peaks can be more securely be linked to abiotic factors, such as paleotemperature. For example, as suggested by Wilberg (2017), the Eocene peak could be related to the Early Eocene Climatic Optimum (Zachos *et al*., 2008), reflecting an increase in diversity (Mannion *et al*., 2015). However, although this relationship was not quantitatively investigated here (i.e. through statistical correlation test), it is difficult to draw more general conclusions, such as that palaeotemperature (or other environmental factor) drives overall patterns of crocodylomorph cranial disparity through time. Similarly, other large-scale investigations of crocodylomorph evolution (such as species diversity and body size evolutionary patterns; Mannion *et al*., 2015; Godoy *et al*., 2019) found more significant influence of abiotic factors only at smaller temporal and phylogenetic scales. This would be consistent with the different biological and physiological characteristics presumed for distinct crocodylomorph subgroups (which range from species highly adapted to a fully-aquatic life to terrestrial and nearly cursorial forms), for which different responses to environmental changes are expected. Accordingly, within the paradigm of adaptive landscapes, overall patterns of phenotypic evolution (such as cranial shape) are more likely to reflect clade-specific adaptations related to the invasion of new adaptive zones (with in turn involve new specific environmental conditions), particularly when analysing large-scale events, across numerous subgroups.

This could also help understanding the nearly continuous decline in crocodylomorph disparity since the Late Cretaceous, which is mainly represented by members of the crown-group Crocodylia. With few exceptions, crocodylians are predominantly semi-aquatic species (Wilberg *et al*., 2019), what could explain their relatively low cranial disparity (Fig. 2c) despite being the most specious crocodylomorph subgroup in my analyses (n = 89). The extinction of other subgroups, which were occupying a wider variety of ecological niches, combined with the presence of predominantly semi-aquatic forms during the Cenozoic, could be a consequence of differential responses to environmental changes (such as global cooling; Zachos *et al*., 2008), which potentially led to a reduced niche availability. Therefore, the current scenario of most modern crocodylian species being piscivorous semi-aquatic animals, within relatively limited variability of body sizes and cranial shapes, could be the result a longstanding pattern of habitat loss in crocodylomorphs, leading to a narrower range of ecologies and morphologies.

## Acknowledgments

I am in debt with my PhD supervisor Richard Butler for support during the development of this study, particularly for encouraging me to go ahead with it as the sole author. I thank Hans Larsson, Trina Du, Felipe Montefeltro, Eric Wilberg, Roger Benson, Andrew Jones, Jeroen Smears, Alexander Beyl and Alan Turner for discussion and methodological assistance at different stages of this study. Access to fossil collections was possible thanks to Lorna Steel (NHMUK), Eliza Howlett (OUMNH), Ronan Allain (MNHN), Rainer Schoch (SMNS), Erin Maxwell (SMNS), Marisa Blume (HLMD), Eberhard Frey (SMNK), Max Langer (LPRP/USP), Sandra Tavares (MPMA), Fabiano Iori (MPMA), Jaime Powell (PVL), Rodrigo Gonzáles (PVL), Martín Ezcurra (MACN), Stella Alvarez (MACN), Alejandro Kramarz (MACN), William Simpson (FMNH), Akiko Shinya (FMNH), Liu Jun (IVPP), Corwin Sullivan (IVPP), Zheng Fang (IVPP), Anna K. Behrensmeyer (USNM), and Amanda Millhouse (USNM). Felipe Montefeltro and Giovanne Cidade also provided photographs of remaining crocodylomorph specimens. This research was supported by the University of Birmingham, Coordenação de Aperfeiçoamento de Pessoal de Nível Superior (CAPES; grant number: 3581-14-4) and the National Science Foundation (grant: NSF DEB 1754596).

## Supporting Information

**Appendix A.** Supplementary methods and results.

**Appendix B.** List of all specimens sampled, including specimen number, group, lifestyle and source of information for landmark digitisation.

**Appendix C.** Landmark dataset (in .tps format), phylogenetic trees (in .tre format), and R script with geometric morphometric, disparity, *SURFACE* and *bayou* analyses.

**Appendix D.** Plots (in .pdf) with all results of disparity-trough-time analyses.

**Appendix E.** Plots (in .pdf) with all *bayou* and *SURFACE* results.

