## Appendix A for "Crocodylomorph cranial shape evolution and its relationship with body size and ecology"

This file contains:

### Supplementary methods

#### *Sampling and data collection*

The dataset published by Wilberg (2017) includes 131 crocodylomorph specimens, most of which were identified to species level. Wilberg (2017) digitised four landmarks on the right side of the skulls of individual specimens (reflecting the left side of the skull when only this was available or it was better preserved or more complete than the right side): “Landmark 1” at the caudalmost point of the skull at the midline (excluding the occipital condyle), “Landmark 2” at the rostralmost point of the snout, “Landmark 3” at the rostralmost point of the orbit margin, and “Landmark 4” at the caudalmost point of the quadrate or quadratojugal (when visible) or the caudolateralmost point of the skull. Apart from these four landmarks, the outlines of skulls were digitised using sliding semilandmarks (Bookstein, 1996; 1997; Adams *et al*. 2004). As described by Wilberg (2017), the outline is traced as a curve extending from “Landmark 1” to “Landmark 2”, on the right side of the skull, and subsequently resampled to contain 60 equidistant semilandmarks (Fig. S1).

However, the dataset made available by Wilberg (2017) was modified from its original version, since it contained semilandmark curves divided into two separate parts (one rostral to and another caudal to the level rostralmost point of the orbit), creating two modules on outline of skulls and potentially exacerbating differences between longirostrine and brevirostrine forms. Therefore, software tpsDIG2 (Rohlf, 2015) was used to manually modify all specimens in the original dataset, resampling the outline curves to contain 60 equidistant semilandmarks.

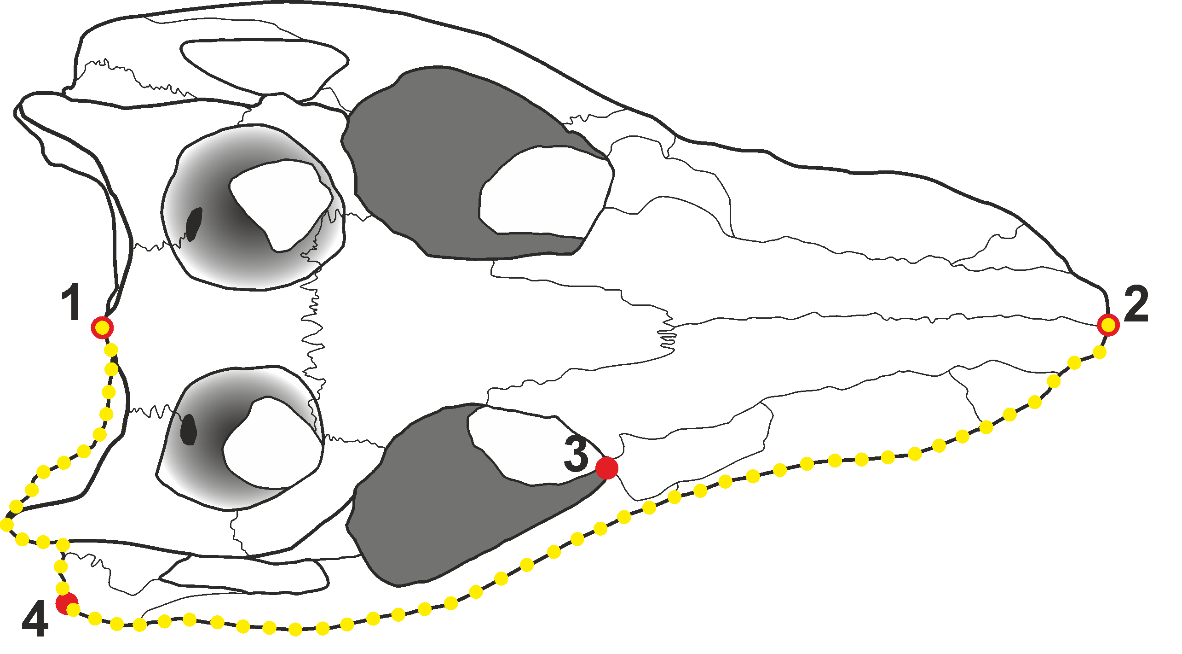

**Figure S1.** Example, using one specimen (*Araripesuchus wegeneri*, MNN GAD19), of the position of the four landmarks (in red) and the 60 equidistant semilandmarks curve (in yellow) used in the present work.

This dataset was then significantly expanded, with the digitisation of landmarks and semilandmarks for 86 new specimens, representing an increase of nearly 65% over the dataset of Wilberg (2017). Non-mesoeucrocodylian crocodylomorphs were particularly underrepresented in Wilberg (2017), with only three taxa present in the original dataset, but 18 in the expanded dataset. However, in general it was possible to expand the taxonomic sample of all crocodylomorph subgroups. From the 131 specimens included in the original dataset, five were removed, including three unnamed specimens (“*Borealosuchus* Tullock specimen”, “*Mecistops* Lothagam specimen”, and “*Prodiplocynodon* Utah specimen”), as well as *Stomatosuchus* (because the fossil material was lost during World War II [Sereno & Larsson, 2009], preventing the inclusion of the taxon in more recent phylogenetic analyses), and *Hamadasuchus* sp. (because first-hand observation allowed me to instead use the holotype specimen of this genus). Following Wilberg (2017), I did not include specimens identified by previous workers as juvenile or sub-adult in the expanded dataset, in order to reduce the effect of ontogeny on cranial shape variation. Furthermore, I also updated the taxonomy, permitting me to identify all specimens at least to species level and to incorporate more recent taxonomic opinions. Thus, some of the specimens treated by Wilberg (2017) as different species were merged (this was mostly the case among thalattosuchians following recent taxonomic work: e.g. Parrilla-Bel *et al*. [2013], Herrera *et al*. [2015], and Foffa *et al*. [2018]). In the cases where multiple specimens are present for a species, averaged values were obtained before the Procrustes alignment.

#### *Phylogenetic framework*

The crocodylomorph informal supertree used in this study is a modified version of the supertree from Godoy *et al*. (2019), which is itself a newer version of the supertrees previously constructed by Bronzati *et al*. (2012; 2015). I used an informal approach to add 20 additional species (by manually modifying the tree using the software Mesquite version 3.51; Maddison & Maddison, 2018), resulting in a final version with 325 species. For the phylogenetic positions of the additional taxa included, I followed recently proposed phylogenetic hypotheses. These included Brochu (2006) for relationships within gavialids, Jouve *et al*. (2015) for tomistomines, Meunier & Larsson (2017) for tethysuchians, Tennant *et al*. (2016) and Schwarz *et al*. (2017) for atoposaurids, and Parrilla-Bel *et al*. (2013), Herrera *et al*. (2015), Fanti *et al*. (2016), Foffa *et al*. (2018), and Ősi *et al*. (2018) for thalattosuchians. Additionally, I followed the phylogenetic hypothesis presented in Wilberg (2017) for some extant crocodylians included in his original landmark dataset.

As mentioned in the main text, for dealing with major phylogenetic uncertainties within Crocodylomorpha, three alternative phylogenetic scenarios were used for the downstream analyses, varying the position of Thalattosuchia, as well as that within the crown-group Crocodylia (i.e. modifying the relative positions of “thoracosaurs”, gavialids and tomistomines). In the first, Thalattosuchia was placed within Neosuchia, and “thoracosaurs” + gavialids formed Gavialoidea (following a more “classic” view; Bronzati *et al*., 2012; 2015). For the second, Thalattosuchia was moved to be the sister taxon of Crocodyliformes (as suggested by some recent hypotheses; e.g. Wilberg, 2015; Wilberg *et al*., 2019). The third topology also places Thalattosuchia sister to Crocodyliformes, but places gavialids and tomistomines together, making “thoracosaurs” non-crocodylian eusuchians by definition (following the results of Lee & Yates [2018]). It is worth mentioning, however, that all the remaining relationships among and within other crocodylomorph subgroups were maintained across these three alternative phylogenetic scenarios.

#### *Geometric morphometric analyses (initial steps)*

The initial steps of the geometric morphometric analyses performed in the present study (i.e. GPA followed by PCA) are, to some extent, different from those applied by Wilberg (2017). The first difference was that during GPA (also known as Procrustes superimposition, Procrustes fit, and GPA-alignment), Wilberg (2017) allowed up to three iterations to occur (with semilandmarks allowed to slide differently in each iteration). As I did not set a maximum number of iterations, this resulted in up to six iterations in my analyses. Wilberg (2017) also used a different software for performing GPA and ordination (tpsRELW; Rohlf, 2015). Furthermore, instead of PCA, Wilberg (2017) used Relative Warp Analysis (RWA) as the ordination method, which should produce equivalent results to PCA, as the author did not weight the variation by bending energy (i.e. α = 0; Zelditch *et al*., 2012; Rohlf, 2015). However, as a sensitivity analysis, I submitted the original landmark data from Wilberg (2017) to the same procedure used for the expanded dataset, in order to estimate the impact of these alternative procedures on the results (i.e. morphospace occupation and PC scores [eigenvalues of each principal component or eigenvector from the PCA]). The resultant morphospace plot (Fig. S2) was very similar to that of Wilberg (2017, Figure 4), indicating that possible discrepancies in the results with the expanded dataset would not be the consequence of the different alignment and ordination methods used.

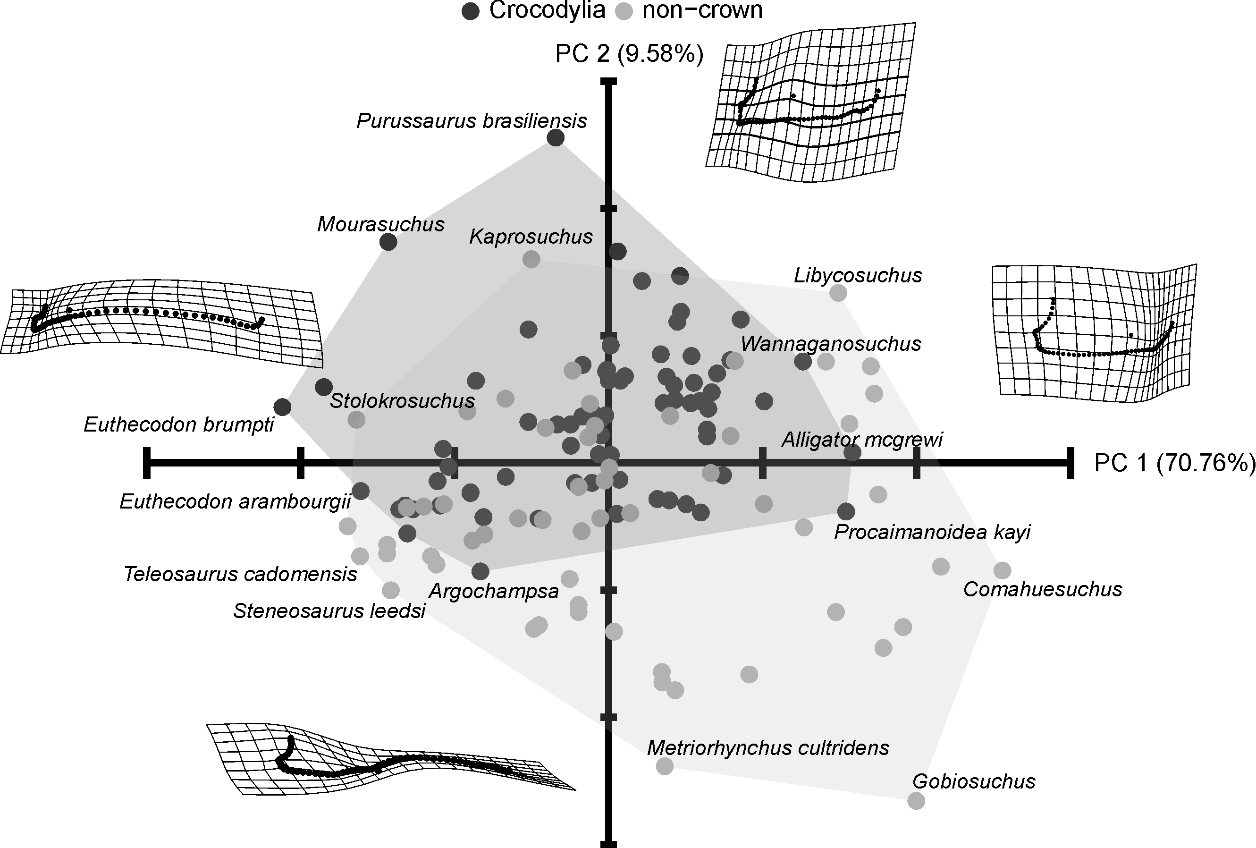

**Figure S2.** Morphospace scatterplot of the first two principal components using only specimens included in the original dataset of Wilberg (2017), with 131 taxa, but using different alignment and ordination methods. Comparative morphospace occupation of crocodylians (crown group) and non-crocodylian crocodylomorphs (non-crown) is shown using convex hulls. Deformation grids illustrate the cranial shape at the extremes of each PC.

#### *Interobserver error*

To quantitatively approach the possible issue of measurement error caused by different observers/operators during the digitisation of landmarks, I used Procrustes ANOVA to compute the amount of variation caused by interobserver error. The Procrustes coordinates of all taxa were used as the response variable in the Procrustes ANOVA formula, and the independent (grouping) variable was a discrete vector indicating whether the landmark data was originally collected by Wilberg (2017) or by me. As recommended by Collyer *et al*. (2015), I used a residual randomization permutation procedure with 10,000 permutations to test the model significance. The percentage of variation driven by interobserver error was then calculated by dividing the sum of squares (SS; see Goodall, 1991) for the term (grouping variable) by the total sum of squares of the model. The results (Table S1) show that interobserver error accounts for only around 1% of the total observed variation, demonstrating the minimal influence of measurement error from different operators during the digitisation of landmarks.

**Table S1.** Procrustes ANOVA results investigating the amount of variation in shape data caused by interobserver error after the standardisation of the landmark digitising protocol. **SS,** sum of squares after 10,000 permutations; **MS,** mean squares; **% of variation,** obtained by dividing the sum of squares of the independent variable (groups) by the total sum of squares; **F,** F-statistic; **p,** *p*-value. *Significant at alpha = 0.05.

| **Total shape variation (all PCs)** | | | | | |
| --- | --- | --- | --- | --- | --- |
| **Effect** | **SS** | **MS** | **% of variation** | **F** | **p** |
| **Interobserver error** | 0.0870 | 0.086985 | 1.606624 | 3.38 | 0.0408* |
| **Residuals** | 5.3272 | 0.025735 |  |  |  |
| **Total** | 5.4142 |  |  |  |  |

#### *npMANOVA*

npMANOVA was used to statistically assesses the differences in the morphospace occupied and cranial shape disparity (sum of variance) by different groups (i.e. taxonomic and ecological groups). In contrast to a parametric MANOVA, npMANOVA does not require the data to be normally distributed, and tests for significant differences on the basis of permutations (Anderson, 2001). Pairwise comparisons between groups were performed, with PC scores of taxa within each group transformed into a Euclidean distance matrix and permuted 10,000 times. I then obtained adjusted *p*-values, to reduce the likelihood of type 1 statistical errors, using the Bonferroni correction (Rice, 1989). It is worth mentioning that, as the Bonferroni correction is considered very conservative, other methods for dealing with multiple comparisons were also applied (such as the “Holm” and the “BH” methods; Holm [1979]; Benjamini & Hochberg [1995]), as a sensitive test. Nevertheless, when different methods provided similar results, only the Bonferroni-corrected *p*-values were reported.

#### *Disparity metric*

There are different methods for quantifying morphological disparity, which for geometric morphometric data correspond to alternative proxies for space occupation (commonly known as disparity metrics or indices; Foote, 1993; 1997; Wills *et al*., 1994; Wills, 2001; Zelditch, *et al*. 2012; Hopkins & Gerber, 2017). Among these, the most frequently used in palaeontological studies are the sums and products of ranges and variances (Wills *et al*., 1994; Guillerme & Cooper, 2018), which usually provide somewhat different results, and have their own limitations (e.g. range metrics are more susceptible to problems arising from uneven sampling, and variance metrics can introduce co-variance between not measured dimensions; Butler *et al*., 2012; Guillerme & Cooper 2018). However, assessing the impact of different metrics on disparity estimating is beyond the scope of this paper. Thus, herein, disparity is defined as the sum of variances expressed for all dimensions (eigenvectors), using PC scores of specimens from all non-null principal components. The sum of variances was selected as it seems to be more robust for measuring disparity through time (see Wills *et al*., 1994; Guillerme & Cooper, 2018), and also because it allows comparisons with the results from Wilberg (2017) as well as other studies (e.g. Stubbs *et al*., 2003; Toljagić & Butler, 2013). Thus, herein, disparity is defined as the sum of variances expressed for all dimensions (eigenvectors), using PC scores of specimens from all non-null principal components.

#### *Sensitivity analyses in disparity-through-time estimation*

As detailed by Guillerme & Cooper (2018), different methods for sub-sampling taxa through time can have important impacts on the results of disparity-through-time studies. For example, using stratigraphic intervals such as stages (a widely employed approach; e.g. Prentice *et al*., 2011; Foth *et al*., 2012; Hughes *et al*., 2013; Stubbs *et al*., 2013; Benton *et al*., 2014; Wilberg, 2017) usually introduces sampling biases, since some short time bins can include very few taxa, leading to large confidence intervals (Guillerme & Cooper, 2018). Alternatively, time bins can be arbitrarily set to represent equal length intervals (e.g. Butler *et al*., 2012; Foth & Joyce, 2016; Foth *et al*., 2016, 2017; Wilberg, 2017), which might diminish the issues of uneven sampling. However, as this approach only allows comparisons of changes in disparity that occur between intervals, it potentially assumes a punctuated model of evolution (i.e. “punctuated equilibrium”, *sensu* Gould & Eldredge, 1977), and prevents the assessment of variation within intervals as a result of gradual evolution (Guillerme & Cooper, 2018). As an alternative to this time binning method, Guillerme & Cooper (2018) recently proposed the “time-slicing” method, which is a phylogeny-based method (i.e. using data from terminal taxa, nodes and branches) and takes into consideration those taxa contemporaneous at specific equidistant points in time (instead of taxa that were present between two points in time), resulting in even sampling. Furthermore, this method allows *a priori* definition of the evolutionary model underlying the changes in disparity.

Therefore, to more rigorously assess the patterns of crocodylomorph cranial shape disparity through time, I used both time sub-sampling methods, with time sub-samples (either time bins or time slices) defined by the number of intervals (i.e. using the same number of time intervals for both methods). In this case, I used 10 and 20 time intervals (i.e. equal-length time bins, in the time binning method, and equally distant specific points in time, in the time-slicing method), to assess the impact of the number of time intervals. In both methods, I used landmark data for both terminal taxa and hypothetical ancestors (obtained with R package *geomorph* [Adams & Otárola-Castillo, 2013], which in turn uses function *fastAnc()* from package *phytools* [Revell, 2012]), and I also allowed taxa to occur in multiple time intervals, by using taxon first and last occurrence dates. PC scores of taxa in each time bin were subjected to bootstrapping (1,000 iterations) to calculate confidence intervals.

For the time-slicing method, I calculated disparity assuming gradual and punctuated models of evolution. For punctuated evolution, selection of values of ancestors or descendants was based on the position of the time slice along the branch, using the score of the ancestor if it is located in the first half of the branch, and the score of the descendant if it is in the second half (i.e. “proximity”; Guillerme & Cooper, 2018). For gradual evolution, a probability function of the distance between the nodes/tip at the ends of the branch and the slice was used for selecting the values (i.e. “gradual splits”; Guillerme & Cooper, 2018).

Finally, for assessing the impact of phylogenetic and temporal uncertainties, I used 10 different time-scaled trees of each alternative topology, which were randomly sampled from the posterior distribution of tip-dating MCMC analyses (i.e. FDB method).

#### *Details of* SURFACE *and* bayou *analyses*

Both *SURFACE* and *bayou* algorithms assume an Ornstein-Uhlenbeck [OU] process (Ingram & Mahler, 2013; Uyeda & Harmon, 2014). OU models include three parameters (theta [*θ*, the trait optimum value], alpha [*α*, the strength of attraction], and sigma squared [*σ*^2^, the rate parameter]), and can be formulated as 𝑑𝑋(𝑡) = 𝛼[𝜃 − 𝑋(𝑡)]𝑑𝑡 + 𝜎𝑑𝐵(𝑡), to expresses the amount of change in trait *X* during the infinitesimal time interval from *t* to *t* + *dt* (Hansen, 1997; Butler & King, 2004; Beaulieu). Both methods allow *θ* to vary across the tree (i.e. allowing for multiple regime shifts), but estimate a single *α* and *σ*^2^ for the entire phylogeny. Accordingly, *SURFACE* and *bayou* can be understood as a non-uniform OU models, whereas the regimes these methods are able to identify across the phylogeny (i.e. by detecting different trait optima) can be interpreted as similar to adaptive zones, within the Simpsonian Adaptive Landscape paradigm (Simpson, 1944; 1953; Stanley, 1973; Hansen, 2012).

*SURFACE* analyses followed the protocol explained in the main text (i.e. backward-phase with pBIC [instead of AICc) and multiple time-scaled trees [given the algorithm sensitivity to phylogenetic and temporal changes]. Furthermore, I discarded shifts with unrealised values of *θ* (i.e. higher or lower than the observed distribution of trait values in my sample). For *bayou* analyses, I used only the MCC trees, but each analysis had 5 MCMC chains of 1,000,000 generations, to ensure convergence. Each chain had starting parameters randomly drawn from the prior distributions to ensure convergence of parameter estimates across chains. I placed normal distributions as priors on *α*, *σ*^2^, *θ* parameters. For number of shifts, a conditional Poisson distribution was used, with a mean equal to 5% of the total number of branches in the tree and a maximum number of shifts equal to half the number of tips. I estimated the effect size, as a predictor of power (Ho & Ané, 2014; Cressler *et al*., 2015; Smaers *et al*., 2017), by calculating the signal-to-noise ratio (SNR), which is obtained with the equation √𝜂𝜙, where 𝜂 is a measure of selection opportunity and 𝜙 is a measure of effect size. Then, I used the SNR for determining a “cut off” value of posterior probabilities of regime shifts (inferring high power when SNR ≫ 1).

### Supplementary results

#### *Principal components*

Principal component analysis (PCA) of the expanded dataset reveal that PC1 (71.89% of the variation) mostly describes variation in snout length (as in Wilberg [2017]), but it also expresses changes in the caudolateral region of the skull, maybe reflecting the addition of many metriorhynchoid thalattosuchians (twice as many as in Wilberg [2017]’s dataset) and some gavialoids (such as *Piscogavialis jugaliperforatus* and *Ikanogavialis gameroi*). PC2 (8.6% of the variation) mostly describes changes in the quadrate condyle and the position of the orbit in relation to the lateral outline of the skull, similarly to Wilberg (2017). All other principal components summed up represent less than 16% of total shape variation (Fig. S3), none of which represents more than 4% (e.g. PC3 and PC4 represent 3.74% and 3.16% respectively).

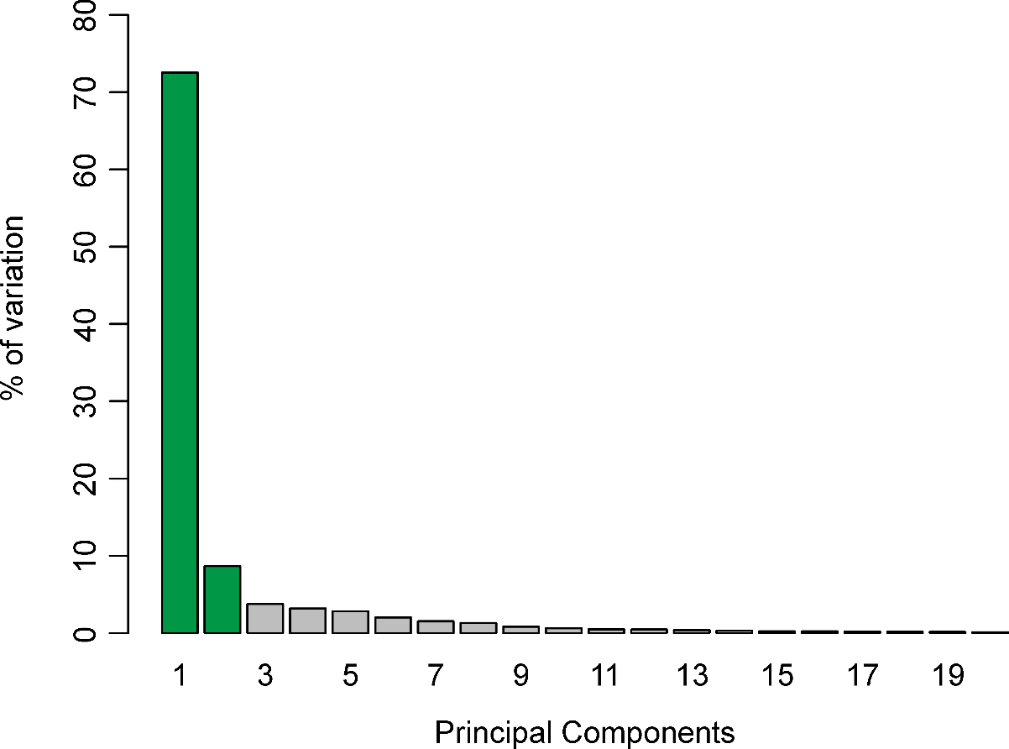

**Figure** **S3.** Percentage of the cranial shape variation expressed by each of the first 20 principal components, using the modified version of the expanded dataset (after the standardisation of the landmark digitising protocol). Shown in green are the principal components (PC1 and PC2) used for morphospace scatterplots in this work.

#### *Morphospace occupation*

The region around the negative end of the PC1 axis (left side of the plot; Fig. S4) shows a concentration of longirostrine species, most of which are gavialoids (such as *Piscogavialis jugaliperforatus* and *Ikanogavialis gameroi*), thalattosuchians (such as *Pelagosaurus typus*, *Steneosaurus bollensis*, and *Teleosaurus cadomensis*), and tethysuchians (such as the “pholidosaurid” *Terminonaris browni* and the dyrosaurid *Rhabdognathus keiniensis*). However, some representatives of other groups are also present in this region, such as the notosuchian *Stolokrosuchus lapparenti* and the crocodyloid *Euthecodon brumpti*. The positive extremity of PC1 (Fig. S4) is dominated by forms with very short rostra, such as the notosuchians *Notosuchus terrestris* and *Comahuesuchus brachybuccalis*, the non-crocodylian neosuchians *Iharkutosuchus makadii* and *Acynodon adriaticus*, as well as the non-mesoeucrocodylian crocodyliform *Gobiosuchus kielanae*. The distribution along the PC2 axis is not as extreme as for PC1 but, in general, taxa at the positive end of the axis (at the top of the plot; Fig. S4) have broader snouts and wider quadrate condyles. This region is occupied by the caimanine alligatoroid *Purussaurus brasiliensis*, as well as some notosuchians (such as *Lorosuchus nodosus* and *Armadillosuchus arrudai*). Notosuchians (such as C*andidodon itapecuruense*) are also observed in the region of negative PC2 scores (at the bottom of the plot; Fig. S4), in addition to many thalattosuchians (such as *Cricosaurus lithographicus*, *Maledictosuchus riclaensis* and *Rhacheosaurus gracilis*).

Compared to crocodylians and notosuchians (see main text for morphospace occupation of these two groups), the morphospaces of thalattosuchians and tethysuchians (Fig. S4) are more restricted to the “longirostrine region” of the scatterplot, and the npMANOVA tests further indicate that their morphospaces do not differ significantly from each other (Table S2). The npMANOVA results also show that the morphospace of non-mesoeucrocodylian crocodylomorphs is not significantly distinct from that of notosuchians, although they exhibit an apparently less disparate morphospace occupation, with nearly all species confined to the region of shorter snouts (Fig. S4). Lastly, even though the morphospace of non-crocodylian neosuchians is, in general, more restricted than that of crocodylians and they do not differ significantly (Fig. S4 and Table S2), some species explore regions of the morphospace not occupied by members of the crown-group (e.g. those of extremely short snouts, such as *Acynodon* and *Iharkutosuchus*).

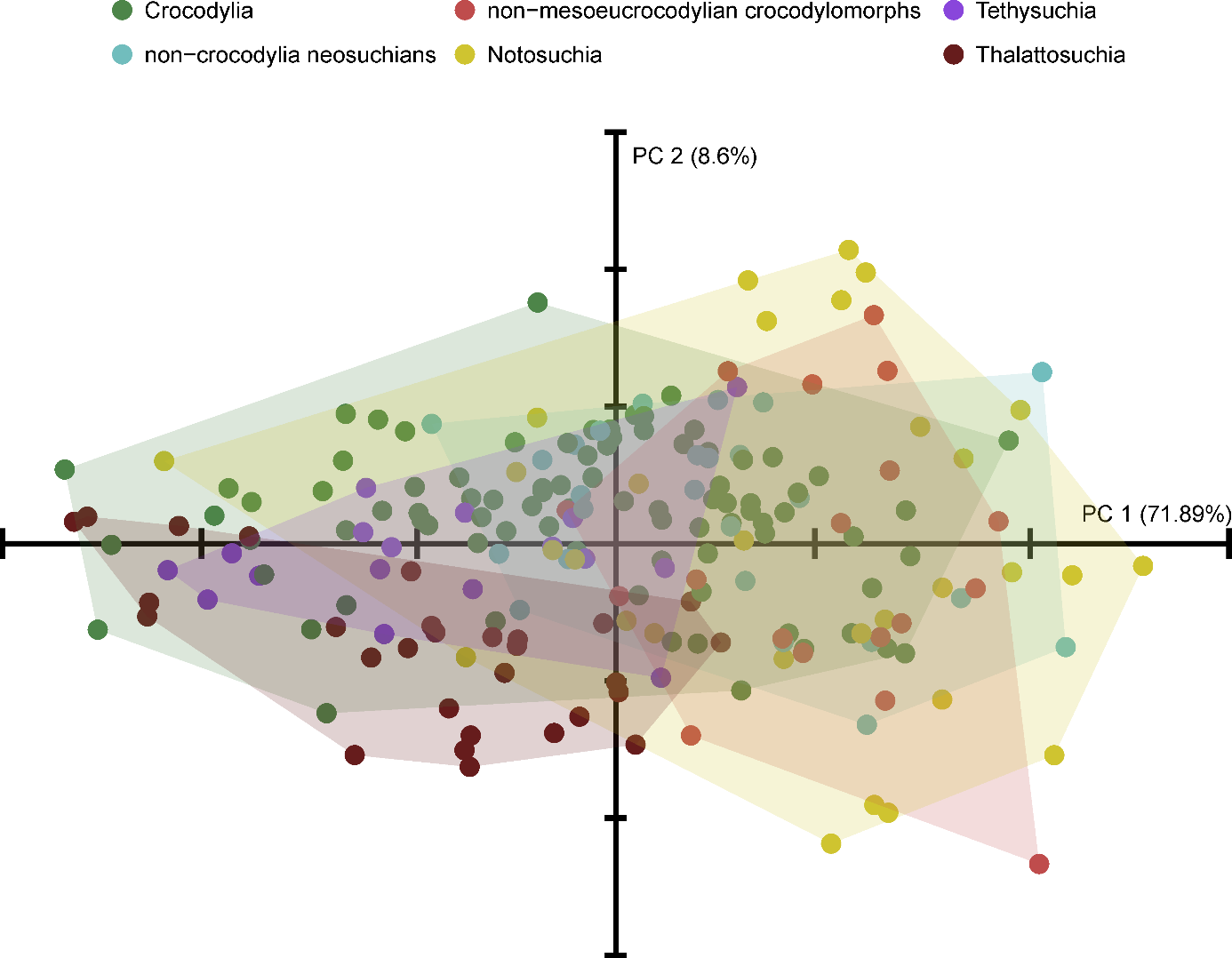

**Figure S4.** Comparative morphospace occupation (PC1 versus PC2) of different crocodylomorph subgroups using convex hulls: Crocodylia (with “thoracosaurs”; n = 89); Notosuchia (n = 30); Thalattosuchia (n = 29); Tethysuchia (n = 18); Non-mesoeucrocodylian crocodylomorphs (without thalattosuchians; n = 18); Non-crocodylian neosuchians (without tethysuchians and thalattosuchians; n = 25).

**Table S2.** npMANOVA results illustrating differences in the morphospace occupied by distinct crocodylomorph subgroups in a pairwise comparison. *Bonferroni-corrected *p*-values (*q*-values) significant at alpha = 0.05.

| Pairwise comparison | *q*-values |
| --- | --- |
| Crocodylia – Notosuchia | 0.0015* |
| Crocodylia – Thalattosuchia | 0.0015* |
| Crocodylia – Tethysuchia | 0.0585 |
| Crocodylia – Non-crocodylian neosuchians | 0.066 |
| Crocodylia – Non-mesoeucrocodylian crocodylomorphs | 0.0015* |
| Notosuchia – Thalattosuchia | 0.0015* |
| Notosuchia – Tethysuchia | 0.0015* |
| Notosuchia – Non-crocodylian neosuchians | 1 |
| Notosuchia – Non-mesoeucrocodylian crocodylomorphs | 1 |
| Thalattosuchia – Tethysuchia | 1 |
| Thalattosuchia – Non-crocodylian neosuchians | 0.0015* |
| Thalattosuchia – Non-mesoeucrocodylian crocodylomorphs | 0.0015* |
| Tethysuchians – Non-crocodylian neosuchians | 0.0015* |
| Tethysuchians – Non-mesoeucrocodylian crocodylomorphs | 0.0015* |
| Non-neosuchians crocodylians – Non-mesoeucrocodylian crocodylomorphs | 0.069 |

#### *Disparity between taxonomic groups*

**Table S3.** npMANOVA results for pairwise comparisons of cranial shape disparity (sum of variances) in different crocodylomorph subgroups (divided into six subsets). *Bonferroni-corrected *p*-values (*q*-values) significant at alpha = 0.05.

| Pairwise comparison | *q*-values |
| --- | --- |
| Crocodylia – Notosuchia | 0.0015* |
| Crocodylia – Thalattosuchia | 0.0015* |
| Crocodylia – Tethysuchia | 0.0015* |
| Crocodylia – Non-crocodylian neosuchians | 0.0015* |
| Crocodylia – Non-mesoeucrocodylian crocodylomorphs | 0.0090* |
| Notosuchia – Thalattosuchia | 0.0015* |
| Notosuchia – Tethysuchia | 0.0015* |
| Notosuchia – Non-crocodylian neosuchians | 0.0015* |
| Notosuchia – Non-mesoeucrocodylian crocodylomorphs | 0.0015* |
| Thalattosuchia – Tethysuchia | 0.0090* |
| Thalattosuchia – Non-crocodylian neosuchians | 0.7919 |
| Thalattosuchia – Non-mesoeucrocodylian crocodylomorphs | 0.6884 |
| Tethysuchians – Non-crocodylian neosuchians | 1 |
| Tethysuchians – Non-mesoeucrocodylian crocodylomorphs | 0.0015* |
| Non-neosuchians crocodylians – Non-mesoeucrocodylian crocodylomorphs | 0.0015* |

As a sensitivity analysis, I also estimated disparity with crocodylians and tethysuchians within Neosuchia, resulting in only four taxonomic subsets (Notosuchia, Thalattosuchia, Neosuchia and non-mesoeucrocodylian crocodylomorphs). The results are presented below (Fig. S5 and Table S5).

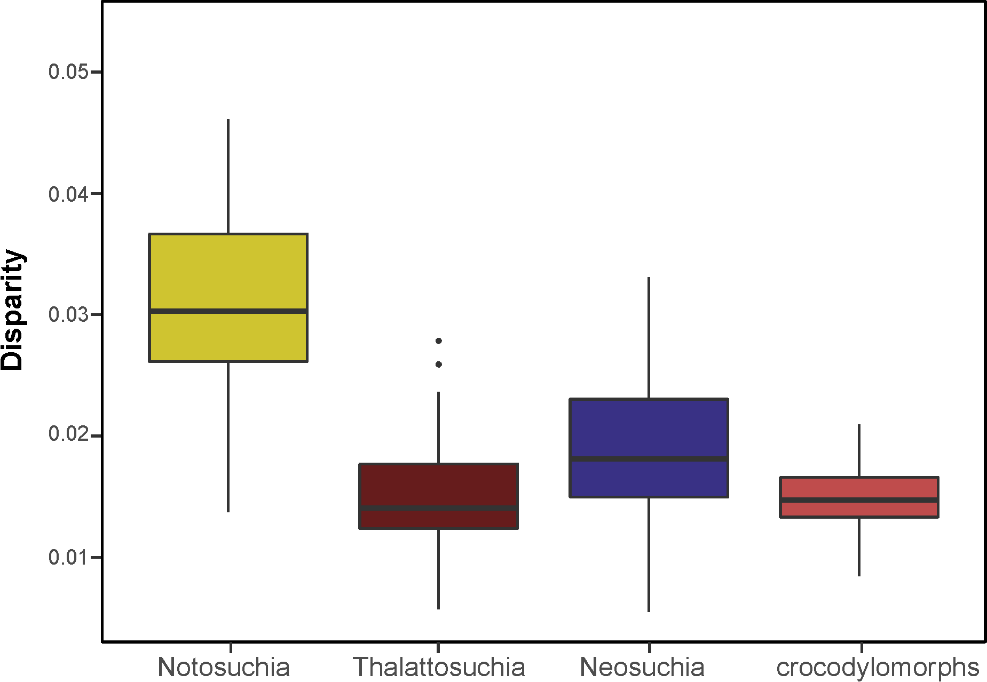

**Figure S5.** Cranial shape disparity (sum of variances) among different crocodylomorph subgroups. Species are divided into four taxonomic subsets (i.e. “Neosuchia” also includes crocodylians and tethysuchians). PC scores of specimens were bootstrapped and rarefied for disparity calculation.

**Table S4.** npMANOVA results for pairwise comparisons of cranial shape disparity (sum of variances) in different crocodylomorph subgroups (divided into four subsets). *Bonferroni-corrected *p*-values (*q*-values) significant at alpha = 0.05.

| Pairwise comparison | *q*-values |
| --- | --- |
| Notosuchia – Neosuchia | 0.0006* |
| Notosuchia – Thalattosuchia | 0.0006* |
| Notosuchia – Non-mesoeucrocodylian crocodylomorphs | 0.0006* |
| Neosuchia – Thalattosuchia | 0.0006* |
| Neosuchia – Non-mesoeucrocodylian crocodylomorphs | 0.0006* |
| Thalattosuchia – Non-mesoeucrocodylian crocodylomorphs | 1 |

#### *Phylomorphospace*

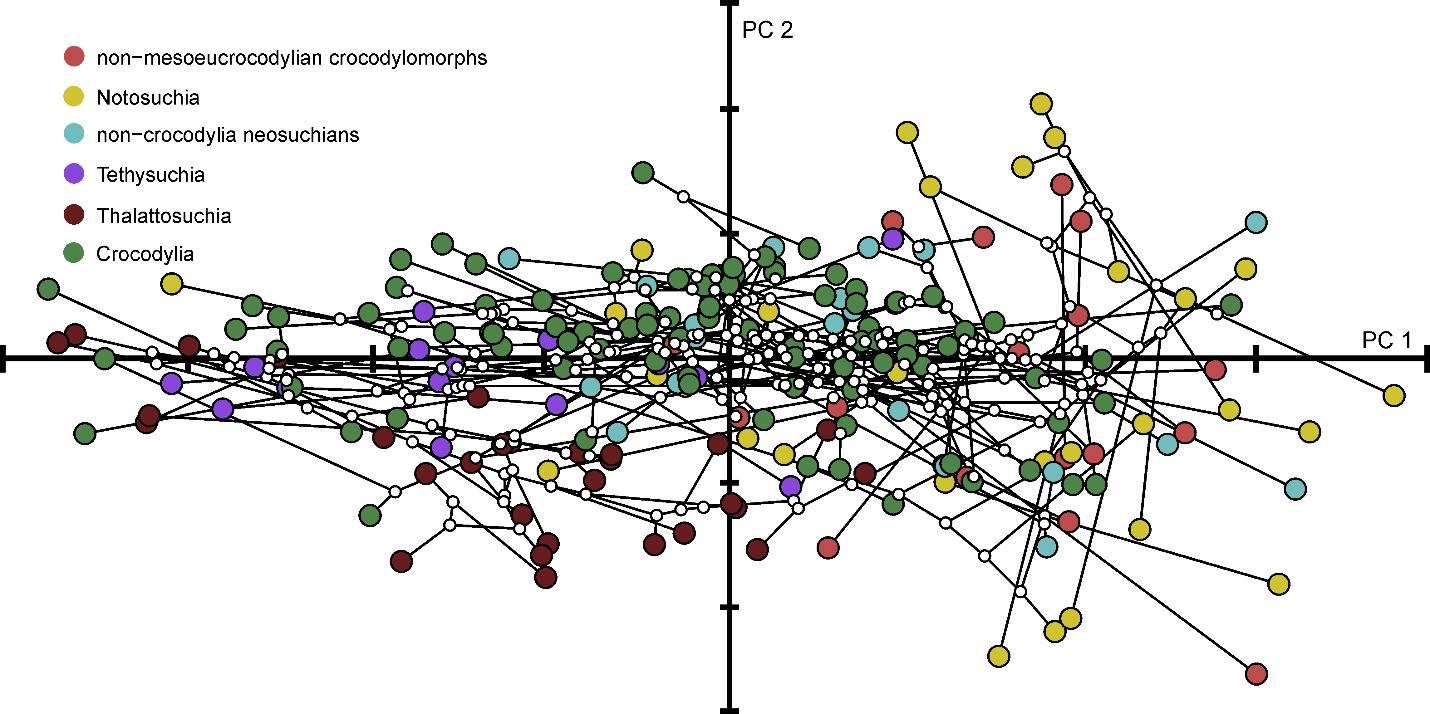

**Figure S6.** Phylomorphospace (i.e. phylogenetic relationships plotted on to PC scores of specimens) of crocodylomorph cranial shape variation (PC1 versus PC2). Circles of different colours represent distinct crocodylomorph subgroups: non-mesoeucrocodylian crocodylomorphs, Notosuchia, non-crocodylian neosuchians, Tethysuchia, Thalattosuchia, and Crocodylia. White circles represent ancestor states (Procrustes coordinates), which were reconstructed using maximum likelihood. Phylogenetic hypothesis from MCC tree, with Thalattosuchia within Neosuchia, but different topologies produce very similar results (i.e. with the multiple intersections of branches).

#### *Disparity-through-time estimation*

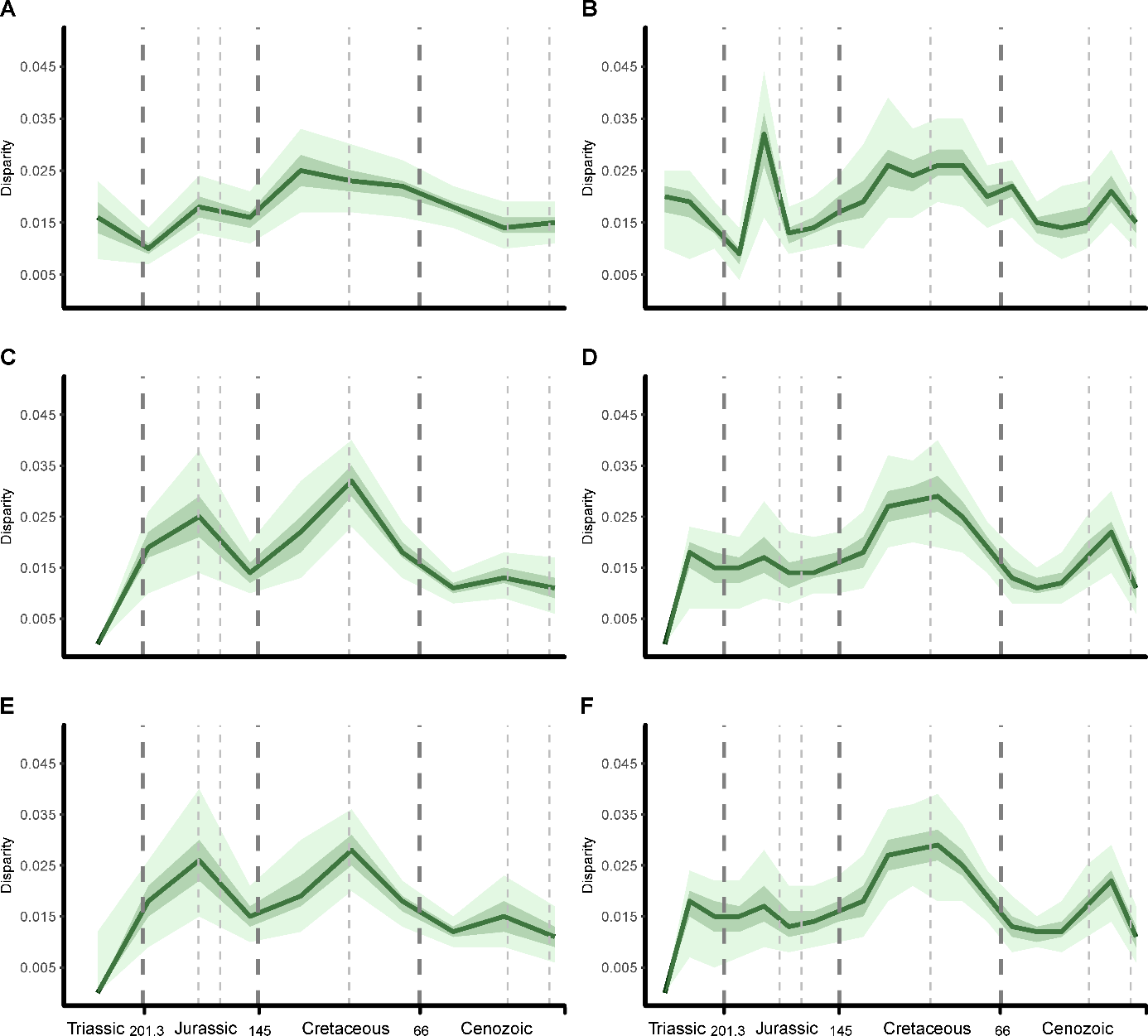

**Figure S7.** Comparison between different time sub-sampling methods used for calculating crocodylomorph disparity through time. All analyses used the same time-scaled tree (tree number 4 with Thalattosuchia within Neosuchia). (a) Time binning method, with 10 equal-length time bins. (b) Time binning method, with 20 equal-length time bins. (c) Time-slicing method, with 10 time slices, assuming punctuated evolution. (d) Time-slicing method, with 20 time slices, assuming punctuated evolution. (e) Time-slicing method, with 10 time slices, assuming gradual evolution. (f) Time-slicing method, with 20 time slices, assuming gradual evolution. The sum of variance is used as the disparity metric. Light and dark green shades represent, respectively, 75% and 97.5% confidence intervals from 1,000 bootstrapping replicates.

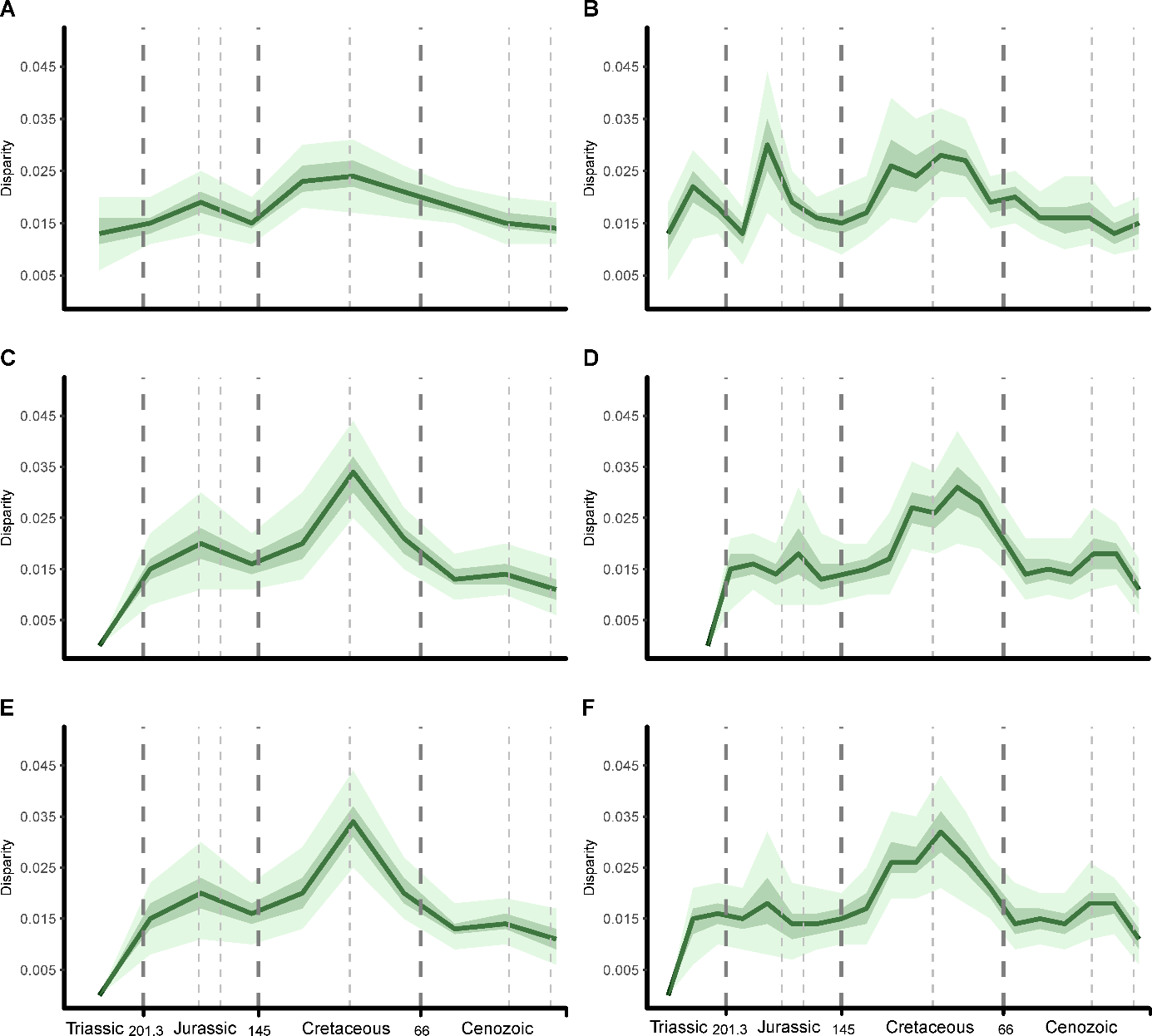

**Figure S8.** Comparison between different time sub-sampling methods used for calculating crocodylomorph disparity through time. All analyses used the same time-scaled tree (tree number 9 with Thalattosuchia sister Crocodyliformes and “thoracosaurs” within Crocodylia). (a) Time binning method, with 10 equal-length time bins. (b) Time binning method, with 20 equal-length time bins. (c) Time-slicing method, with 10 time slices, assuming punctuated evolution. (d) Time-slicing method, with 20 time slices, assuming punctuated evolution. (e) Time-slicing method, with 10 time slices, assuming gradual evolution. (f) Time-slicing method, with 20 time slices, assuming gradual evolution. The sum of variance is used as the disparity metric. Light and dark green shades represent, respectively, 75% and 97.5% confidence intervals from 1,000 bootstrapping replicates.

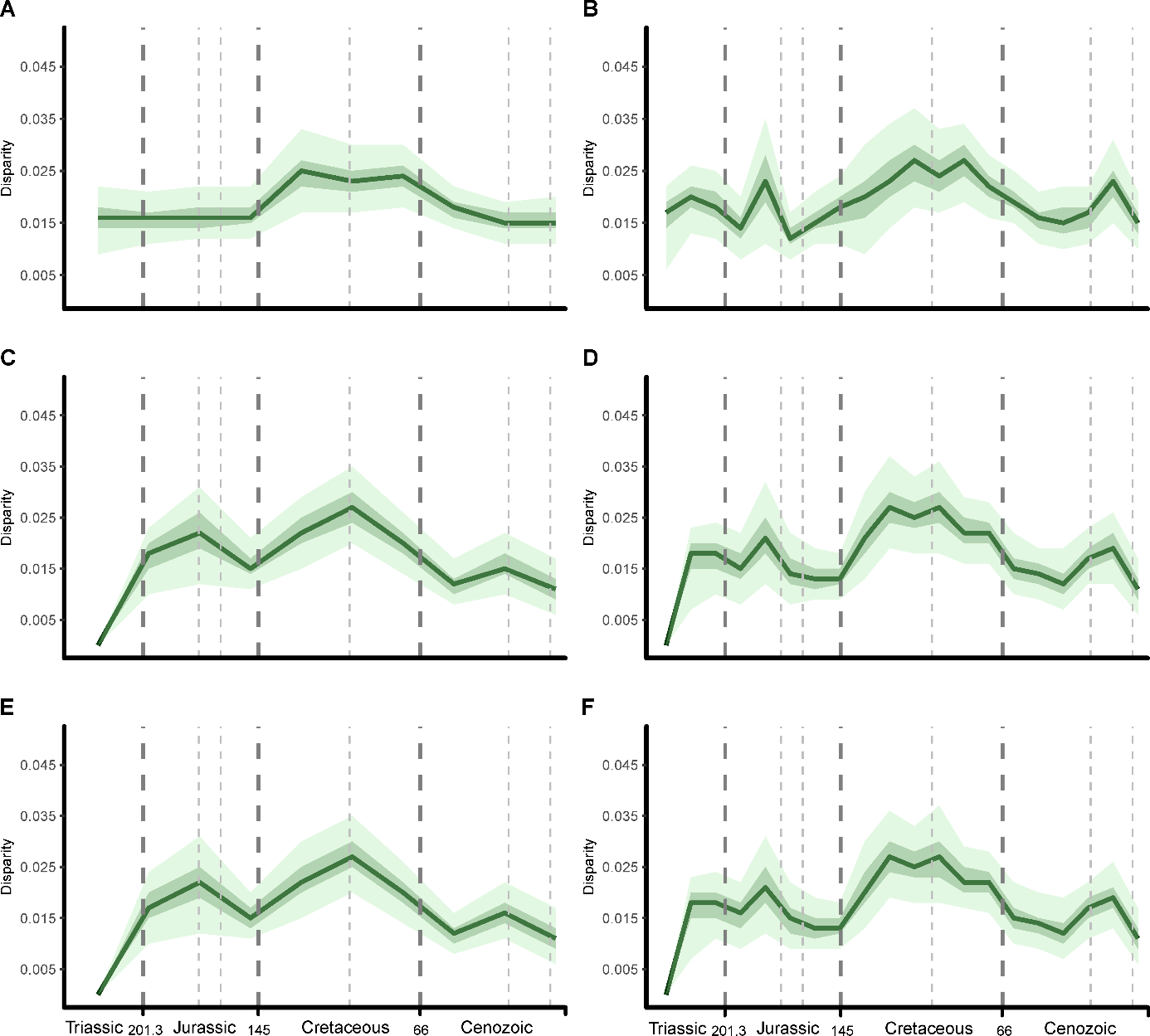

**Figure S9.** Comparison between different time sub-sampling methods used for calculating crocodylomorph disparity through time. All analyses used the same time-scaled tree (tree number 8 with Thalattosuchia sister Crocodyliformes and “thoracosaurs” outside Crocodylia). (a) Time binning method, with 10 equal-length time bins. (b) Time binning method, with 20 equal-length time bins. (c) Time-slicing method, with 10 time slices, assuming punctuated evolution. (d) Time-slicing method, with 20 time slices, assuming punctuated evolution. (e) Time-slicing method, with 10 time slices, assuming gradual evolution. (f) Time-slicing method, with 20 time slices, assuming gradual evolution. The sum of variance is used as the disparity metric. Light and dark green shades represent, respectively, 75% and 97.5% confidence intervals from 1,000 bootstrapping replicates.

#### *Size-shape relationship*

**Table S5.** Results of regressions of shape data (i.e. PC1 scores) on body size data (i.e. an independent dataset of log-transformed dorsal cranial length measurements, from Godoy *et al*. 2019). Possible correlation was analysed using ordinary least squares (OLS) and phylogenetic generalised least squares (PGLS) regressions, this latter one with three phylogenetic scenarios (with alternative positions of Thalattosuchia and gavialids; topologies 1, 2 and 3). In all regressions, n = 160. *Significant at alpha = 0.05.

| **OLS** |  |  |  |
| --- | --- | --- | --- |
| **R^2^** | **Intercept** | **Slope** | **AIC** |
| 0.5457 | -0.71132 | 0.28371* (<0.00001) | -300.9822 |
| **PGLS (topology 1)** |  |  |  |
| **R^2^** | **Intercept** | **Slope** | **AIC** |
| 0.2059 | -0.493520 | 0.188777* (<0.00001) | -314.9359 |
| **PGLS (topology 2)** |  |  |  |
| **R^2^** | **Intercept** | **Slope** | **AIC** |
| 0.2971 | -0.554858 | 0.221742* (<0.00001) | -333.1843 |
| **PGLS (topology 3)** |  |  |  |
| **R^2^** | **Intercept** | **Slope** | **AIC** |
| 0.2717 | -0.541737 | 0.214824* (<0.00001) | -308.0961 |

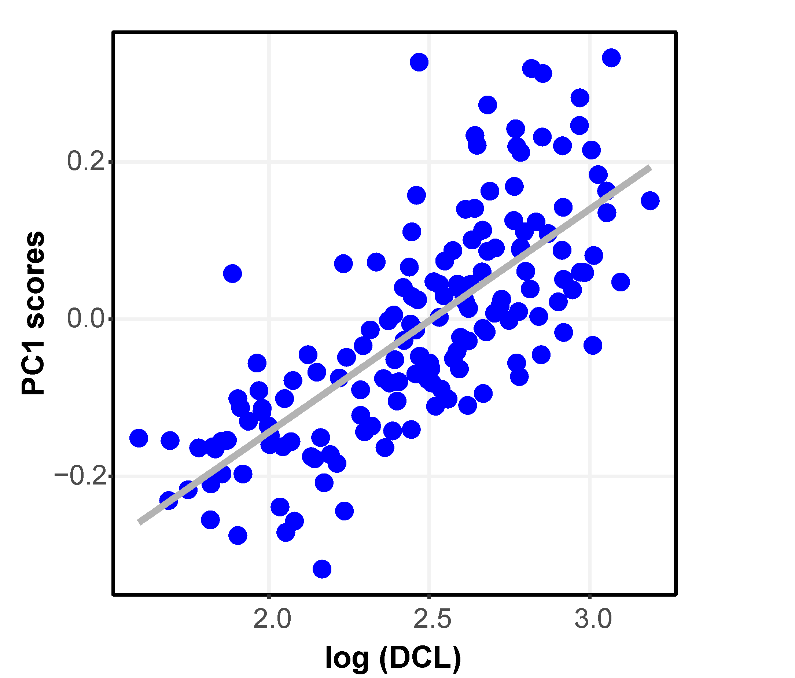

**Fig. S10.** The relationship between cranial shape (represented by PC1 scores) and body size (log-transformed dorsal cranial length [DCL], in millimeters, from Godoy *et al*. 2019) in crocodylomorphs, using ordinary least squares (OLS) regression. Regression results (statistics) shown in Table S5.

#### *Disparity between ecological groups*

**Table S6.** npMANOVA results (pairwise comparisons) of cranial shape disparity (sum of variances) of crocodylomorphs assessed in different lifestyle categories. *Bonferroni-corrected *p*-values (*q*-values) significant at alpha = 0.05.

| Pairwise comparison | *q*-values |
| --- | --- |
| Aquatic – Semi-aquatic | 0.096 |
| Aquatic – Terrestrial | 0.0003* |
| Semi-aquatic – Terrestrial | 0.0003* |

### Institutional abbreviations

**MNN,** Muséum National du Niger, Niamey, République de Niger.
